# A systems approach discovers the role and characteristics of seven LysR type transcription factors in *Escherichia coli*

**DOI:** 10.1101/2021.12.22.473864

**Authors:** Irina A. Rodionova, Ye Gao, Jonathan Monk, Nicholas Wong, Richard Szubin, Hyun Gyu Lim, Dmitry A. Rodionov, Zhongge Zhang, Milton H. Saier, Bernhard O. Palsson

**Affiliations:** Department of Bioengineering, University of California San Diego, La Jolla, CA 92093-0116, USA; Division of Biological Sciences, University of California San Diego, La Jolla, CA 92093-0116, USA; Sanford-Burnham-Prebys Medical Discovery Institute, La Jolla, CA 92037, USA; A.A. Kharkevich Institute for Information Transmission Problems, Russian Academy of Sciences, Moscow, Russia; Department of Pediatrics, University of California San Diego, La Jolla, CA 92093, USA; Novo Nordisk Foundation Center for Biosustainability, Technical University of Denmark, Lyngby 2800, Denmark

**Keywords:** LysR-type transcription factors, DNA-binding sites, ChIP-exo, phenotype microarray, regulon elucidation

## Abstract

Although *Escherichia coli* K-12 strains represent perhaps the best known model bacteria, we do not know the identity or functions of all of their transcription factors (TFs). It is now possible to systematically discover the physiological function of TFs in *E. coli* BW25113 using a set of synergistic methods; including ChIP-exo, growth phenotyping, conserved gene clustering, and transcriptome analysis. Among 47 LysR-type TFs (LTFs) found on the *E. coli* K-12 genome, many regulate nitrogen source utilization or amino acid metabolism. However, 19 LTFs remain unknown. In this study, we elucidated the regulation of seven of these 19 LTFs: YbdO, YbeF, YgfI, YiaU, YneJ, YcaN, YbhD. We show that: 1) YbdO regulation has an effect on bacterial growth at low pH with citrate supplementation. YbdO is a repressor of the *ybdNM* operon and is implicated in the regulation of citrate lyase genes (*citCDEFG*); 2) YgfI activates the *dhaKLM* operon that encodes the phosphotransferase system involved in glycerol and dihydroxyacetone utilization; 3) YiaU regulates the *yiaT* gene encoding an outer membrane protein, and *waaPSBOJYZU* operon is also important in determining cell density at the stationary phase; 4) YneJ, re-named here as PtrR, directly regulates the expression of the succinate-semialdehyde dehydrogenase, Sad (also known as YneI), and is a predicted regulator of *fnrS* (a small RNA molecule). PtrR is important for bacterial growth in the presence of L-glutamate and putrescine as nitrogen sources; and 5) YbhD and YcaN regulate adjacent y-genes on the genome and YbeF is involved in flagella gene regulation. We have thus established the functions for four LTFs and identified the target genes for three LTFs.

**IMPORTANCE:** The reconstruction of the transcriptional regulatory network (TRN) is important for gram-negative bacteria such as *E. coli*. LysR-type TFs are abundant in Enterobacteria, but many LTF functions still remain unknown. Here we report putative functions of uncharacterized TFs based on multi-omics data related to L-threonine, L-glutamate, and putrescine utilization. Amino acids (AAs) and polyamines are important sources of nitrogen for many microorganisms, but the increase in one amino acid or putrescine concentration in a minimal medium also induces stress. Although polyamine metabolism has been studied, the TRN that controls the putrescine (Ptr) and AA utilization at minimal medium conditions are still poorly understood. The function of previously uncharacterized transcriptional regulators YbdO, YgfI, and YneJ (PtrR) were identified in *Escherichia coli*. PtrR is important for Ptr and L-glutamate utilization, while YgfI transcriptional regulation was found to be important for growth on L-threonine and glycerol as a carbon source.

## INTRODUCTION

Recently, the roles of uncharacterized LysR-type transcription factors (LTFs) have been identified via multiple approaches, including transcriptome analysis of uncharacterized TF (yTF)-deleted mutants (machine-learning-based), gene clustering, and the detection of DNA-binding sites ^1^. The predicted yTF targets annotated as transporters and enzymes define the TF physiological role. A combination of the knowledge from EcoCyc, Fitness Browser, and iModulonDB with TF DNA-binding data provided hypotheses for the putative physiological functions of yTFs under different growth conditions ^2^. Indeed, as an example, the function of one TF, PunR, was recently discovered to be an activator for *punC*, purine transporter.

The RNA-seq data for the TF deletion mutants under specific growth conditions provide information about transcription affected by mutation (Fig. 1). In *Escherichia coli*, LTFs are regulators for amino acids (AAs), purine, and dicarboxylate metabolism, nitrogen assimilation (NAC), antibiotic resistance, and virulence. LTF regulatory proteins protect 50-60 bp regions with TA-rich regulatory binding sites and activation binding sites (ABS) and regulate different metabolic pathways. LTFs are known to regulate conserved gene clusters that are adjacent to the genes encoding the regulator^3^, but additionally, LTF autoregulation is a common property. As one of the largest families of HTH-type regulators, LTFs contain an N-terminal helix-turn-helix DNA-binding domain and C-terminal co-inducer binding domain (Fig. S1). Given the broad conservation of LTFs, it is possible that they regulate a wide variety of target genes with diverse physiological functions using common regulatory features.

**Figure 1.**
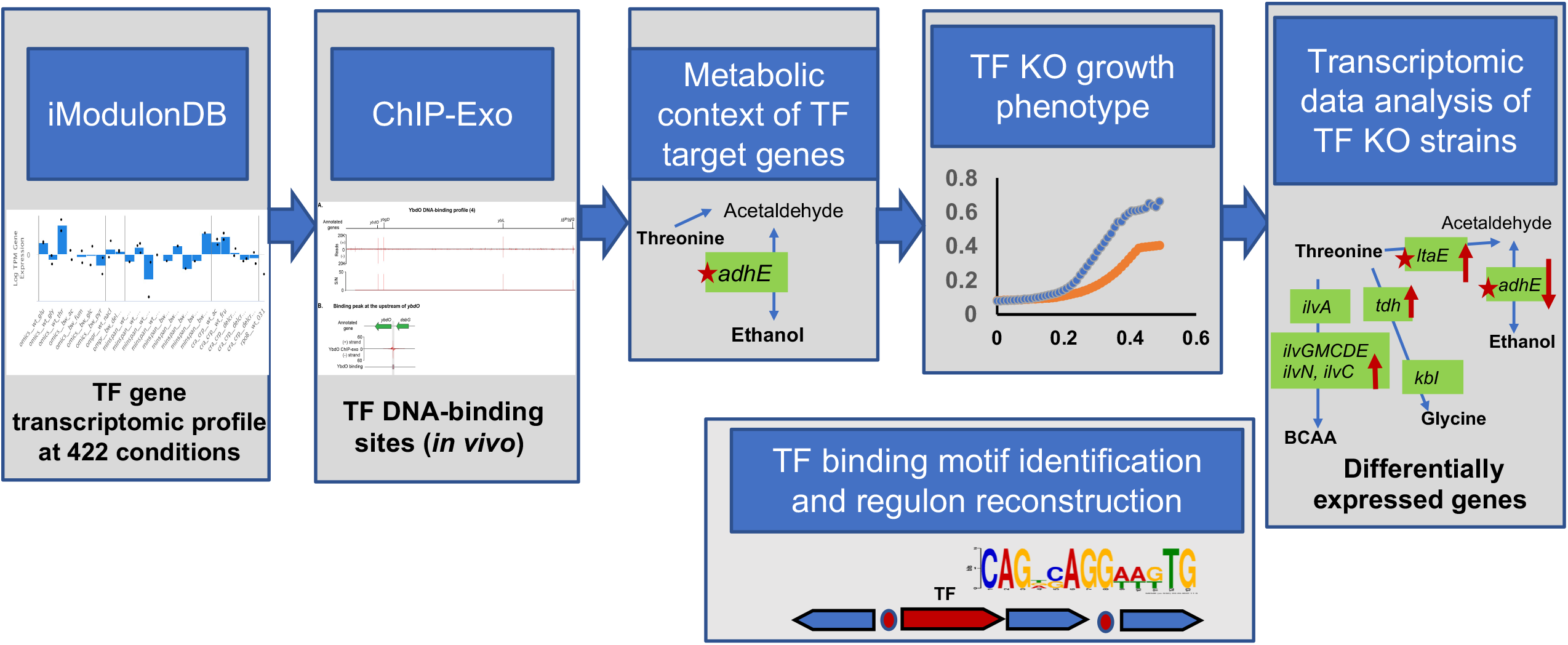
Systems approach for the prediction of transcription factor’s function. A systematic approach to identify LysR family unknown transcriptional factors physiological function in *E. coli*.

LTFs account for 16.7% (47 out of 280) of the total number of transcription factors in *Escherichia coli* K-12 ^4^. Out of the 47 LTFs, 26 (AbgR, AllS, ArgP, Cbl, CynR, CysB, Dan, DmlR, DsdC, GcvA, HcaR, HdfR, IlvY, LeuO, LrhA, LysR, MetR, Nac, NhaR, OxyR, PerR, PgrR, QseA, QseD, TdcA, XapR) have known functions and the majority were shown to regulate adjacent genes ^5^. However, there are still many uncharacterized TFs belonging to the LysR-type family, which require further studies to determine their regulatory functions. This effort is also important for the reconstruction of the transcriptional regulatory network (TRN).

*Escherichia coli* is a representative of the commensal mammalian intestinal microbiota and is the best characterized model gram-negative bacterium. Nutrient starvation conditions are important for the gut microbiome bacterial community as they cause stress, activating different survival mechanisms ^6^, and TRNs rewire the metabolism under different conditions. iModulonDB is a collection of *E. coli* MG1655 transcriptomics data for different growth medium and stress conditions ^7^. Machine-learning based ICA data analysis (iModulonDB), RNA-seq data for the TF knockout, and ChIP-exo data (*in vivo* DNA-binding sites) are useful resources for LTF characterization. Recently, the ChIP-exo results for verified uncharacterized TFs in *E. coli* MG1655 was published ^1^. The gene expression profiling for LTFs under multiple growth conditions in iModulonDB provides important information for predicting the growth conditions for the yTFs ^7^.

We performed systems analysis for the prediction of LTF functional role using data combined from previously published chromatin immunoprecipitation with exonuclease treatment (ChIP-exo) detected LTF DNA-binding sites, transcriptome data from iModulonDB (Fig. 1), RNA-seq analysis for LTF deletion mutants (experimental data), and genome cluster analysis using the bioinformatics tools (Fig.1). We generated RNA-seq data for seven LTF mutants (*ycaN, ygiF, ybeF, ybdO, ybhD, yiaU*, and *yneJ*), analyzed the conserved genome clustering with LTF genes, and detected conserved genes that were differentially expressed in the LTF knockout. Accordingly, we generated a hypothesis about the possible regulatory targets of three LTFs and their physiological functions in response to the high L-threonine concentration in minimal medium (Fig. 1). The LTF regulation in response to the imbalance of Thr and AA catabolism for energy has been studied. The LTF deletion effect on the regulation of biochemical pathways as the biosynthesis of L-serine, L-glycine, and branched-chain AAs from Thr and metabolism of glycerol, ethanol, and formate (Fig. 1) had been investigated in *E. coli* BW25113.

For the detailed analysis, we investigated the effect of the YneJ transcriptional response in minimal medium under nitrogen starvation conditions. The *yneJ-sad* cluster is conserved in enterobacterial genomes, and Sad is involved in putrescine utilization as nitrogen source ^8^. Many bacterial species, including *E. coli*, can simultaneously utilize L-glutamate (Glu) and the polyamine putrescine (Ptr) under carbon/nitrogen starvation conditions. Glu is also essential for tetrahydrofolate polyglutamylation ^9^. Ptr is important for bacterial growth and for efficient DNA replication, transcription, and translation ^10,11^ and plays an important role in maintaining compact conformations of negatively charged nucleic acids ^12^. Ptr is also involved in multiple antibiotic resistance mechanisms under stress conditions ^13^. The *puuA, puuD, puuE*, and *puuP* genes in *E. coli* are induced by Ptr and regulated at the transcriptional level by the Ptr-responsive repressor PuuR ^14,15^. The expression of *sad* (*yneI*) is induced by the addition of Ptr to the medium ^16^; however, it is not regulated by PuuR, and a transcriptional regulator for *sad* had not been described before this work.

Here, using a transcriptomic analysis systems approach, we identify the LysR-type TFs, important for Thr utilization in minimal medium, and YneJ (PtrR), that directly controls *sad* gene expression in response to GABA, an intermediate of the Ptr utilization pathway. The predicted PtrR binding site in the *sad* promoter region was confirmed via an *in vitro* binding assay with the purified PtrR protein and using the ChIP-exo assay. We further compared the whole genome transcriptional response of the *ptrR* knockout and wild type *E. coli* strains to carbon/nitrogen starvation in the presence of Ptr and L-glutamate using RNA-Seq analyses, and the PtrR DNA-binding site was predicted for *fnrS*, encoding small regulatory RNA. The physiological roles of PtrR in the FnrS regulation of anaerobic response and in antibiotic resistance are discussed.

## RESULTS

### An integrated systems-approach uncovers LysR-type transcription factors in *E. coli* K-12

Previously, we generated a list of candidate transcriptional factors (TFs) from uncharacterized genes (“y-genes”) using a homology-based algorithm ^17^. Among these TFs, it was predicted that YeiE, YfeR, YidZ, YafC, YahB, YbdO, YbeF, YbhD, YcaN, YdcI, YdhB, YeeY YfiE, YgfI, YhaJ, YhjC, YiaU, YneJ, and YnfL belong to the LTF based on Hidden Markov Model. YdhB (re-named PunR) function had been shown to be an activator for the purine transporter, PunC ^1,2^. We further chose 7 uncharacterized LysR-type TFs (yLTFs) from the yLTFs, which have transcriptional responses (increased mRNAs) to the presence of a L-threonine (Thr) in M9 medium (iModulonDB, PRECISE2). L-threonine is an important source for L-serine, L-glycine, branched-chain amino acid biosynthesis, and formate, which is important for anaerobic respiration. To elucidate roles of each yLTF and genome-wide target genes, we performed gene expression profiling via RNA-Seq and ChIP-exo detection of the LTF DNA-binding and growth phenotyping. The overall workflow is shown in Fig. 1.

### YbdO and YbeF regulatory effects are involved in citrate utilization

The YbdO ChIP-exo result detected the peak for DNA-binding upstream of *ybdO-ybdNM* (Fig. 2A, B). Further, gene expression profiling showed that *ybdO* deletion leads to strong upregulation of the *ybdNM* operon, suggesting direct regulation of *ybdO*-*ybdNM* and the conserved citrate lyase encoding genes *citCDEF* (Fig. 2C). Therefore, this result confirmed autoregulation. The *ybdO* and *ybdNM* operons are conserved adjacent genes in Enterobacteriaceae (Fig. 2D) and the ybdN promoter was predicted to be regulated by FNR. From PRECISE2, upregulation of *citC* was detected when *E. coli* MG1655 strain was grown in M9 medium supplemented with Thr. The YbdM homologue has been predicted as yTF (Uniprot). The DNA-binding site upstream of the *ybdO* gene was predicted by the phylogenetic footprinting method (Fig. S2). The gene expression profiling results show that YbdO represses *ybdO-ybdNM* and likely indirectly regulates *citCDEGF* (Table 1), as the predicted palindrome binding sequence was not found upstream of *citCDEFG*. To test the regulation of the citrate lyase encoding gene cluster (*citEFG*), the phenotype for growth in the M9 medium supplemented with citrate at low pH (pH 6.5) showed an effect of *ybdO* deletion on citrate utilization (Fig.2E-F), but no difference for the growth in M9 medium supplemented with citrate was found at pH 7.5. Additionally, *ybdO* mutation decreases the *E. coli* BW25113 fitness phenotype for motility in LB medium, and affects growth using glycolate as carbon source (Fitness Browser, fit.genomics.lbl.gov). The data analytic method ICA has evolved independently modulated sets of genes (iModulons) from bacterial transcriptomes (iModulons). The FlhDC iModulon includes FlhDC regulated genes that were upregulated in the *ybeF* and *ybdO* deletion mutants (Fig. S4). YbdO and YbeF are paralogs (identity 30%), presented in conserved gene clusters in Enterobacterial genomes with citrate lyase operon *cit* (*citCDEFXGT*, Fig. 2C, Fig. S3). *citC* transcription is upregulated in the presence of Thr in M9 medium (iModulonDB, PRECISE2). Interestingly the transcriptomic analysis of the *ybdO* and *ybeF* deletion mutants showed that the expression level of *lrhA* was substantially decreased (−4.3 -fold and -3.5 -fold, respectively, Table 1, Supplementary data), likely indicating that YbeF and YbdO has affected *lrhA* regulation (YbdO/YbeF DNA-binding was not detected/predicted). LrhA is an LTF repressor for *flhDC*, flagellar biosynthesis genes responsible for motility and chemotaxis ^18^. The deletion of *ybeF* showed downregulation of *lrhA* and upregulation of *flhC*. The iModulon FlhDC contain flagella encoding genes, regulated by FlhDC (iModulon “FlhDC”, Fig. S4) was substantially upregulated in *ybeF* mutant strain.

**Figure 2.**
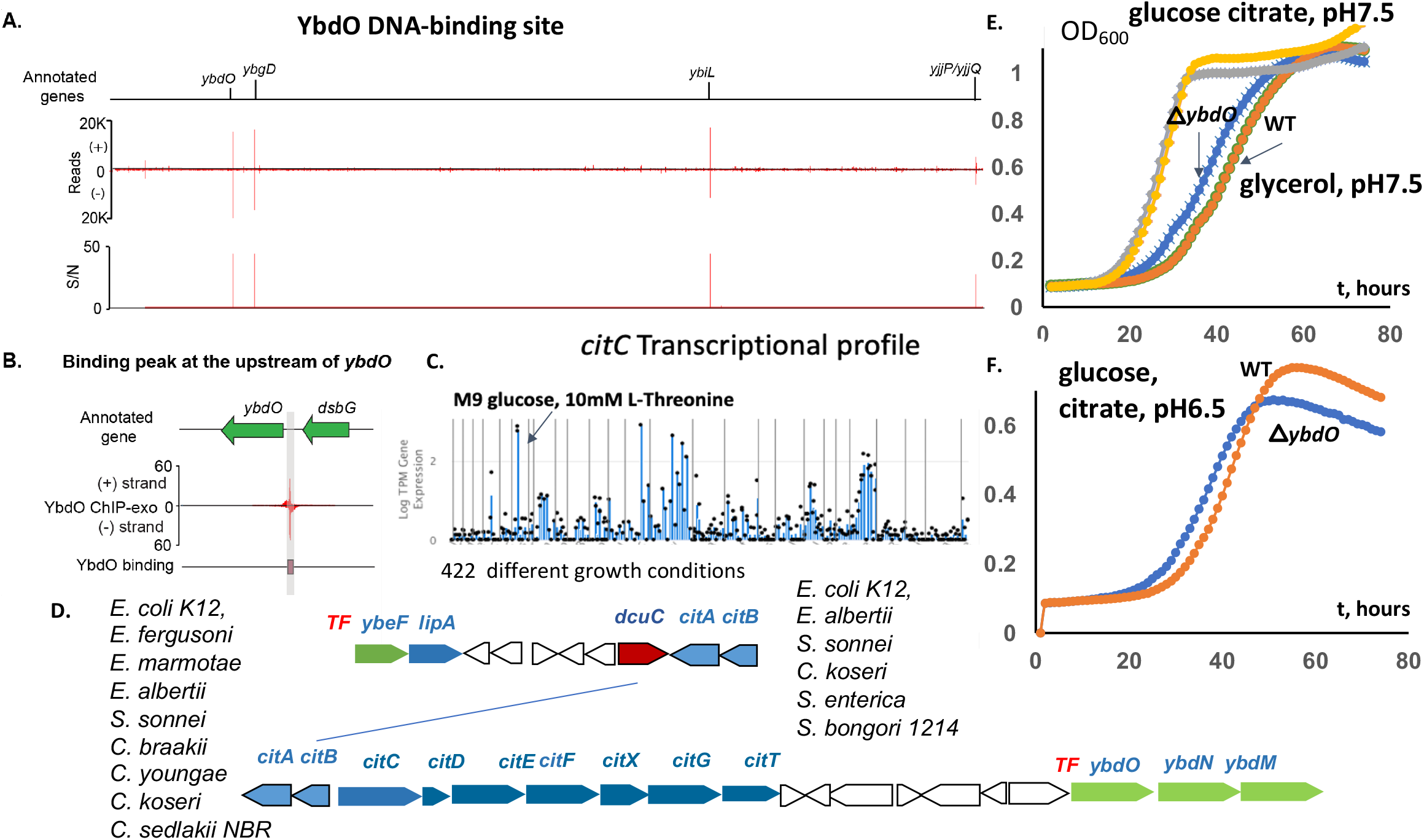
The systems approach for YbdO and YbeF transcriptional factors function prediction. **(A)** The genome-wide binding of YbdO. **(B)** The zoom-in of the binding site at the promoter of the gene *ybdO*. **(C)** Transcriptomic data for *citC* (citrate transporter) **(D)** *citCDEFXGT* (citrate lyase encoding) genes clustering with *ybdO-ybdMN*, and *ybeF*, the analysis across the closely related bacteria. *dcuC*-anaerobic dicarboxylate transporter, *lipA*-lipoyl synthase **(E-F)** The growth measurement of the *ybdO* mutant (orange and grey line) compared to the wild type BW25113 strain of *E. coli* (blue and yellow line). Growth was measured in 96-well plates with glucose (or glycerol) as the carbon source, supplemented by 30 mM citrate at pH 6.5.

### YcaN and YbhD regulators mutant characterization

The *ybhD and ycaN* mutant strains and WT were collected at the late-exponential phase after growth in the M9 medium supplemented with Thr. The conserved gene cluster *ycaN*-*ycaK*-*ycaM* with adjacent *ycaC* and *ycaD* genes was detected in the *Escherichia coli* K12 and *Shigella boydii* genomes. We noticed that in the *ycaN* deletion mutant strain differentially expressed genes *ycaC* (downregulated) and *ycaD* (upregulated) are genes adjacent to *ycaN*. It is interesting that *ycaK, ycaC*, and *ycaD* are predicted to be regulated by Nac (belonging to LTF), as Nac function is nitrogen assimilation, and *ycaC* and *ycaD* are probably nitrogen assimilation function related (EcoCyc). In *ycaN* deletion mutant highly DEGs are: the *tnaAB* genes, encoding L-tryptophanase and a tryptophan transporter, L-arginine degradation, *astCADBE*, an autoinducer-2 transport system (*lsrABCD, lsrR*), and the HTH-type transcriptional regulator, *galS*, the genes were strongly downregulated (Supplementary Data 1). The genes encoding amino acid metabolism (L-valine biosynthesis (IlvB, IlvN), threonine dehydrogenase (Tdh), transcriptional activator (TdcA), and glycolate utilization pathway (GlcC, GlcD, GlcF)) were additionally downregulated in the *ycaN* mutant.

The *ybhD* gene is divergently oriented with respect to the conserved Enterobacterial gene cluster *ybhHI* and the putative hydrolase gene, *ybhJ* (Fig. S5A). YbhH is a 4-oxalomesaconate tautomerase homologous protein, and YbhI is a putative tricarboxylate transporter, homologous to 2-oxoglutarate/malate translocator, (id. 35%) (iModulonDB) (Fig. S5B). The YbhH encoding gene was strongly upregulated in a *ybhD* deletion mutant, as shown by transcriptomic analysis. *ybhH* and *ybhI* transcription is regulated by Nac (EcoCyc, iModulonDB) and the functional relation to nitrogen assimilation is suggested. The *ybhD* deletion strain growth phenotype in the M9 minimal medium with glycerol (carbon source), supplemented by L-malate was detected (Fig. S6).

### YgfI regulation and glycerol and formate utilization

ChIP-exo assays previously detected YgfI binding upstream of the *dhaKLM* operon encoding the dihydroxyacetone phosphotransferase (DHAK) (Fig.3A-B). The transcriptional activation of DhaKLM is important for glycerol utilization and M9 supplemented by Thr or L-tryptophan (Fig. 3B) (iModulonDB). DHAK in the *E. coli* MG1655 strain is involved in glycerol utilization (Uniprot). Accordingly, we decided to test the effect of the *ygfI* deletion on the growth on glucose (Fig. 3C-D) and glycerol (Fig. 3E). The resulting deficiency in growth on glycerol is potentially explained by *dhaKLM* as well as *pflB, hycBCD, hycEFG*, and *adhB* transcriptional YgfI activation, as shown by RNA-seq (Table 2), and the YgfI binding site was predicted upstream of those genes (Table S1). The DEGs detected by RNA-seq showed substantial downregulation of formate fermentation related genes (Fig. 3F) encoding pyruvate-formate lyase (*pflB*), fumarate reductase (*frdABCD)*, formate hydrogenlyase (*hycBCD, hycEFG)*, and the regulator (*hycA*), as well as hydrogenase encoding genes (*hybABC, hybEF*, and *hybO*) and the gene encoding protein involved in maturation of all hydrogenases isozymes (*hypB*) (Fig. 3F) and the *dhaKLM* operon (Table 2, Supplementary Data). *ygfI* deletion had little effect on growth using glucose as the carbon source (Fig. 3C) aerobically, but the *ygfI* strain had a low growth rate microaerobically without Thr supplement in minimal medium (Fig. S7).

**Figure 3.**
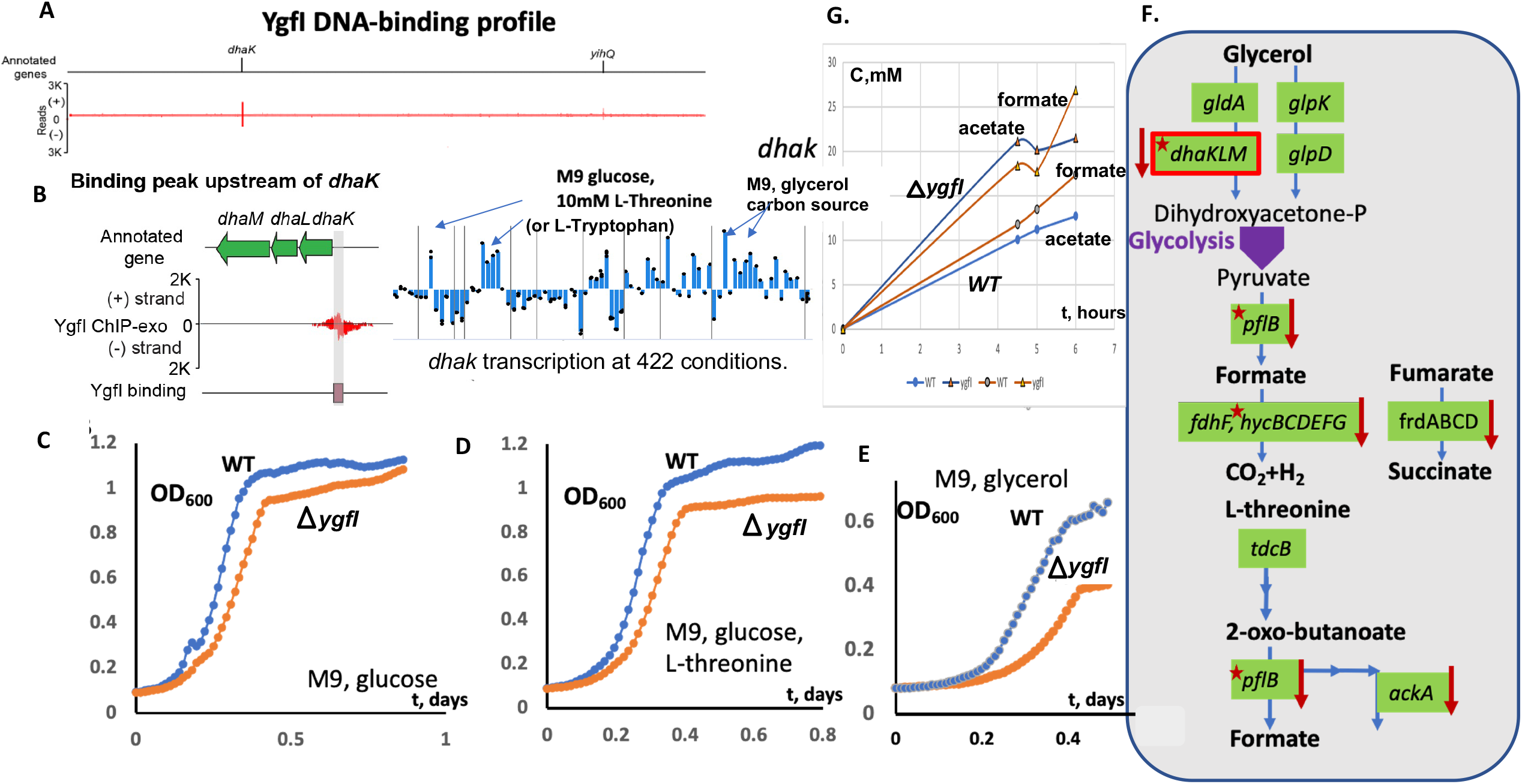
The systems approach for YgfI transcriptional factor function prediction. **(A-B)** The genome-wide binding of YgfI and the zoom-in for the *dhaK* binding site. Transcriptomic profile for the YgfI regulated *dhaK* gene. **(D-E)** The YgfI function was assessed based on the growth phenotype under different conditions. The growth measurement of the *ygfI* mutant (orange line) compared to the wild type BW25113 strain of *E. coli* was measured in 96-well plates in M9 medium or the same medium supplemented with 7 mM L-threonine **(C)** or M9 medium with glycerol as the carbon source **(F)** The glycerol utilization pathway – glycerol kinase (GK, *glpK*), glycerol-3phosphate dehydrogenase (G3PDH, *glpD*), glycerol dehydrogenase (GlyDH, *gldA*), dihydroacetone kinase (DHAK, *dhaKLM*). RNA-seq differentially expressed genes are marked by a red arrow. Transcriptomic data analysis for the *ygfI* mutant compared to the wild type BW25113 strain of *E. coli* shows the glycerol and L-threonine utilization pathway genes (the DEGs are marked by arrows, predicted YgfI binding site marked by red stars) regulation. **(G)** The concentration (mM) for the formate (orange line) and acetate (blue line) produced by *E. coli* BW25113 WT (circles) and *ygfI* deletion mutant (triangles) strains in M9 glucose medium supplemented by 7 mM Thr (microaerobic conditions).

The detected YgfI binding upstream of *dhaKLM* and the glycerol growth phenotype for the *ygfI* mutant confirm the YgfI-dependent transcriptional activation of DHAK. We suggest re-naming YgfI to DhfA - dihydroxyacteone/formate utilization activator. The fermentation products during microaerobic growth on M9 glucose were detected in the Thr supplemented M9 medium. The analysis evolved the higher efflux of formate and acetate for the *ygfI* mutant strain (Fig. 3G), suggesting that YgfI is important for formate utilization as an energy source.

### YiaU regulatory network and yiaU mutant growth phenotype

ChIP-exo results show the YiaU binding for the regulation of *waaP* (encoding lipopolysaccharide (LPS) biosynthesis glycero-D-manno-heptose kinase) (Fig. 4A). The results suggest that YiaU is important for LPS biosynthesis at specific conditions and the majority of DEGs encode the proteins involved in cell wall/membrane/envelope biogenesis, carbohydrate transport, energy production, and amino acid metabolism (Fig. 4B). Accordingly, gene expression for the *yiaU* mutant was lower for the genes from the operon *waaPSBOJYZU*, suggesting that YiaU is the LPS biosynthesis operon activator; previously the regulator for *waa* operon was not known (ecocyc.org). Additionally, ChIP-exo detected binding for the genes *yjiT* and adenine transporter, *adeP*, suggests transcriptional regulation, and they were detected as DEGs for the *yiaU* mutant (Fig. 4D). The difference for the growth in exponential/log phase for the *yiaU* mutant strain had not been detected (Fig. 4C), although YiaU regulation had affected final OD_600_ (stationary phase) in M9 glucose medium. The phenotype (log phase) for the mutant in M9 with 0.3 M NaCl added was additionally detected (Fig. 4C).

**Figure 4.**
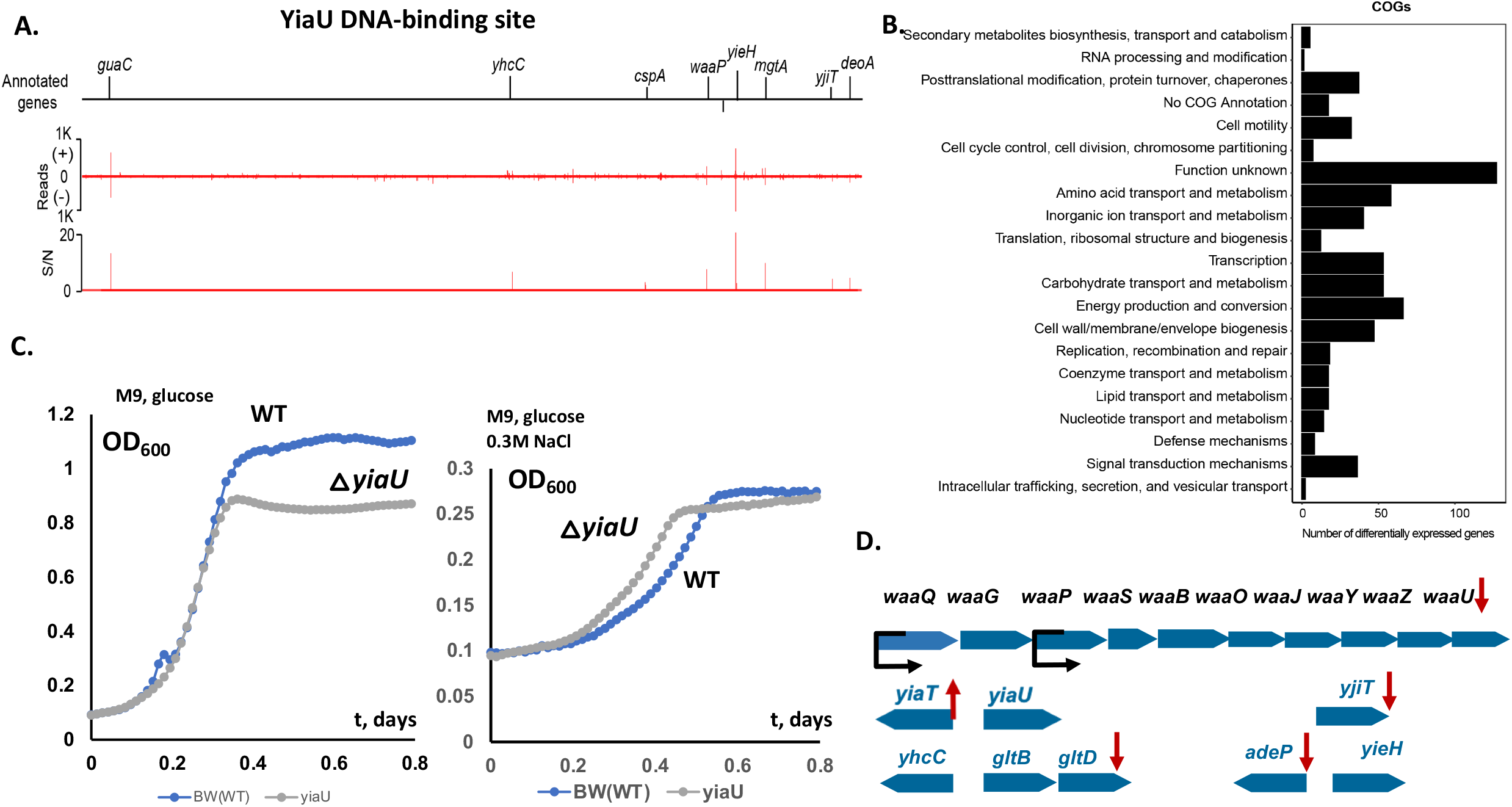
The systems approach for the function of the transcription factor YiaU. **(A)**. The genome-wide binding of YiaU across the genome. **(B)**. Clusters of Orthologous Groups (COGs) were enriched among the differentially expressed genes between the wild type BW25113 and *yiaU* mutant strains. **(C)**. The growth measurement of the wild type and *yiaU* mutant strains under different conditions. Left panel: M9 glucose with L-threonine. Right panel: M9 glucose with L-threonine and 0.3 M NaCl. **(D)**. Predicted structure and YiaU regulation of *waa* operon, *yiaT, yjiT*, and *adeP* in *E*.*coli* BW25113. The differentially expressed genes are shown by a red arrow. The *waa* operon promoters are shown by a black arrow.

The LysR family representatives are known to regulate adjacent genes, and *yiaU-yiaT* are conserved in the bacterial genomes (Fig. 4D). The RNA-seq results for the *yiaU* deletion mutant strain showed upregulation of the *yiaT* gene, encoding a predicted outer membrane protein membrane anchor for the surface display for the proteins, homologue of MipA. MipA is an MltA (murein-degrading enzyme) interacting protein.

### YneJ (PtrR) regulatory effects for *sad* and *fnrS* transcription and putrescine utilization

Two distinct Ptr utilization pathways are known for *E. coli* (Fig. 5A). The first is catalyzed by the PuuA, PuuB, PuuC, and PuuD enzymes encoded by the *puuP, puuA, puuDR, puuCB, puuE* gene cluster and involves degradation of Ptr to γ-aminobutyric acid (GABA) via γ-glutamylated intermediates. The alternative pathway of Ptr degradation to GABA consists of PatA (Ptr aminotransferase) and PatD (γ-aminobutyraldehyde dehydrogenase) ^19^. The PuuABCDE pathway is essential for Ptr utilization in *E. coli* using PuuP as the major Ptr transporter ^20^. GABA is further utilized by two alternative 4-aminobutyrate aminotransferases (GABA-AT) encoded by *gabT* and *puuE*, and also two succinate semialdehyde dehydrogenases (SSADH) encoded by *gabD* and *sad* ^21,22^.

**Figure 5.**
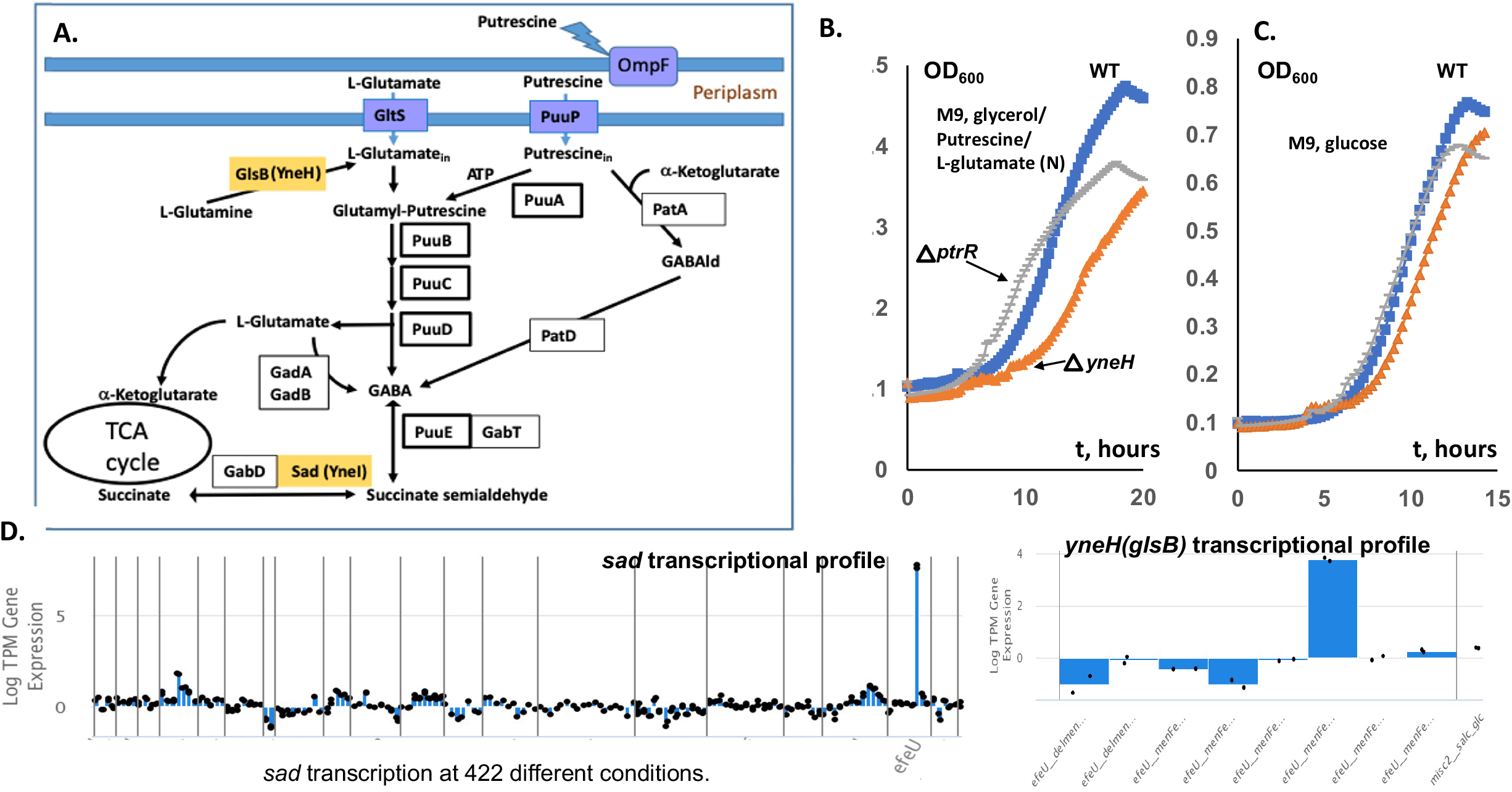
The experimental validation of the transcription factor YneJ (PtrR). (**A)** Overview of two alternative pathways of putrescine utilization in *E. coli*. PtrR-regulated genes are shown in yellow boxes. Transporters are shown in blue. Abbreviations: GABAld – gamma-aminobutyraldehyde, PatA - putrescine aminotransferase, PatD - gamma-aminobutyraldehyde dehydrogenase, YneI (Sad) - succinate-semialdehyde dehydrogenase, YneH (GlsB) - glutaminase. GadAB-two glutamate decarboxylase isoforms, GabD, Sad (YneI)-succinate- semialdehyde dehydrogenase, GlsB(YneH), glutaminase. (**B)** The growth measurement of the *ptrR, yneH* mutants compared to the wild type BW25113 strain of *E. coli*. The cell cultures were grown in M9 medium with 10 mM Glu/10 mM Ptr, as nitrogen sources and 0.4% glycerol (v/v) as the primary carbon source or (**C)** M9 glucose medium. (**D)** Transcriptomic data for *Sad (yneJ)* and *glsB (yneH)* at different growth conditions for *E. coli* MG1655 strain and adapted MG1655 derivatives (iModulonDB, PRECISE2). The activation of the *sad* and *glsB* promoter at the *ptrR* mutant strain was detected.

We decided to characterize in detail the YneJ (re-named PtrR) by analyzing the PtrR ChIP-exo detected binding sites ^1^. The *ptrR* gene is located in a conserved gene cluster with the divergently transcribed *sad* (*yneI)* gene, which encodes succinate semialdehyde dehydrogenase and *yneH* (glsB) glutaminase (Fig. 6A and 6C), upregulated in the evolved *yneJ* mutant strain (iModulonDB) ^23^. To identify and characterize DNA binding sites of PtrR in the *E. coli* genome we utilized a combined bioinformatics and experimental approach. First, we applied a comparative genomic approach of phylogenetic footprinting ^22^ to predict putative PtrR-binding sites in the common intergenic region of the *ptrR* and *sad* genes (Fig. 6C, Fig. S8). The *ptrR*/*sad* genes are conserved in several taxonomic groups including *Escherichia/Salmonella/Shigella, Citrobacter, Enterobacter*, and *Klebsiella*, as well as in *Pseudomonas* spp. In *E. coli* and closely related enterobacteria the *sad* gene belongs to the putative *sad-yneH* gene cluster, while in *Enterobacter* and *Citrobacter* the orthologous genes include an additional gene encoding the methyl-accepting chemotaxis protein I (serine chemoreceptor protein, Mcp) (Fig. 6C). The multiple sequence alignment of *ptrR*/*sad* upstream regions from *E. coli* and closely related enterobacteria (termed Group 1 species) contains a conserved 15-bp palindromic sequence with consensus TTCACnAATnGAGAA downstream predicted sigma-E dependent promoter (Fig. 6A). We also analyzed upstream regions of *ptrR* orthologs in other enterobacterial genomes (Group 2 species), where the *sad* gene ortholog is absent and *ptrR* is co-localized with an uncharacterized MFS-family transporter gene. Further, we identified two conserved DNA sites with similar consensus sequences located in their common intergenic region (Fig. 6C).

**Fig. 6.**
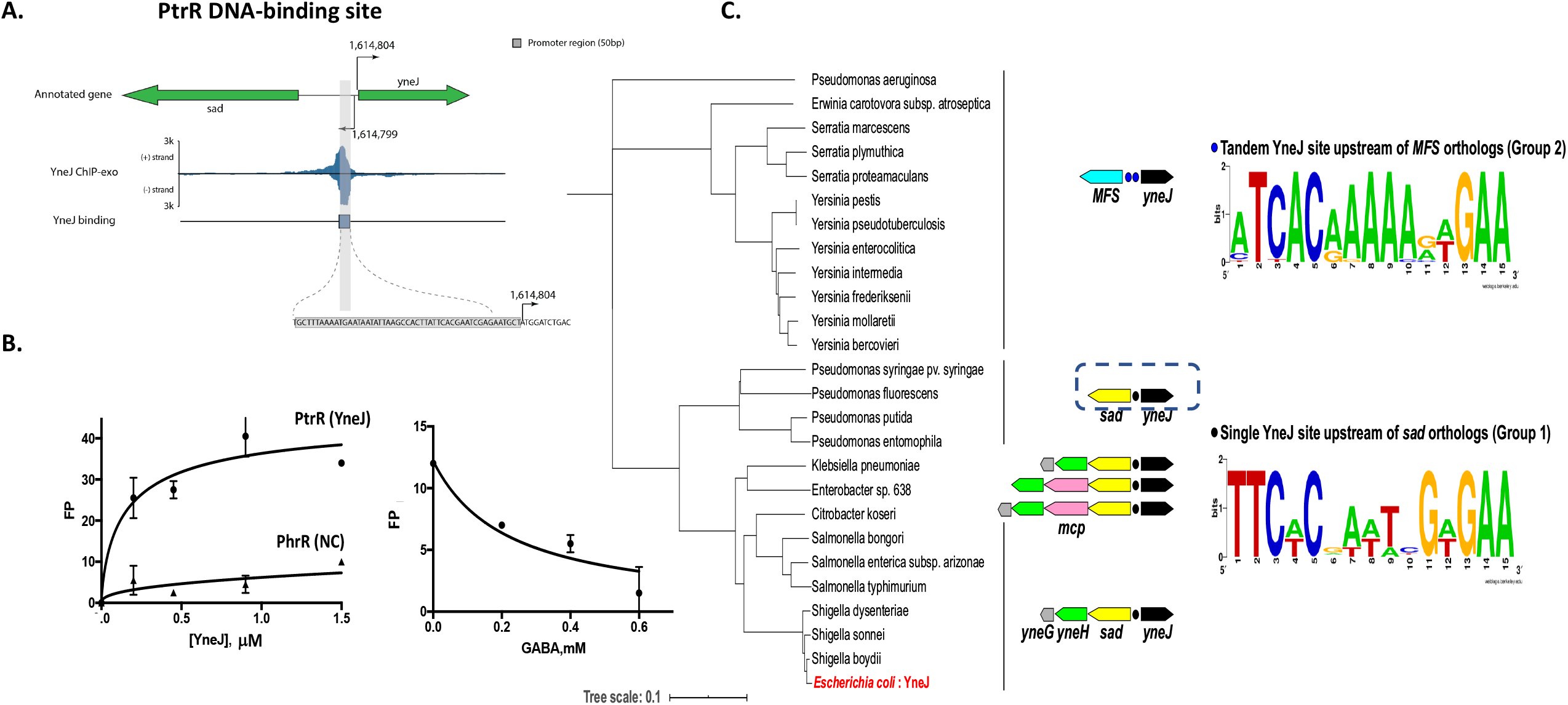
The systems approach for the function of the transcription factor PtrR (YneJ). **(A)** The zoom-in of the PtrR-binding site at the promoter region of the *ptrR* (*yneJ*) and *sad* genes. Location of sigma-H and sigma-E promoters is from the EcoCyc database. **(B)** Fluorescent polarization assay of PtrR binding to the predicted operator site at *sad* gene. PhrR protein from *Halomonas* was used as a negative control. **(C)** The phylogenetic tree of PtrR orthologous proteins and predicted PtrR-binding motifs in *E. coli* and related genomes of Enterobacteria and *Pseudomonas* spp. The maximum likelihood phylogenetic tree was constructed using RAxML. The distinct genomic context of *ptrR* genes from two major tree branches (groups of PtrR orthologs) is shown by arrows with the following colors: black (*ptrR* regulator), yellow (*sad* for succinate semialdehyde dehydrogenase), green (*yneH* for glutaminase), pink (*mcp* for methyl-accepting chemotaxis protein), and blue (mfs for putative MFS-family transporter), while the predicted PtrR-binding sites are shown by black dots. Sequence logos of predicted DNA binding sites of PtrR from each of the two groups of analyzed species were constructed using WebLogo 2.0.

We further confirmed the identified putative PtrR-binding site upstream of the *sad* genes in *E. coli* and conducted genome-wide mapping of other PtrR-binding sites using the ChIP-exo method. To identify *in vivo* PtrR binding sites, *E. coli* was grown under glucose as the carbon source in the M9 minimal media. A total of nine PtrR-binding sites were detected in these experiments. PtrR binds in the promoter regions of the *fhuC, moeB, dhaK, fnrS, uxaB, gltS*/*xanP*, and *ptrR/sad* genes, as well as in the intragenic region of the *yhiQ* and *ydiF* genes. The experimentally identified 50-bp PtrR-binding region at *sad/ptrR* genes contains the conserved palindromic DNA motif identified via phylogenetic footprinting (Fig. S8). Comparison of this DNA motif with eight other regions containing experimentally mapped PtrR-binding regions did not reveal significant sequence similarity except for the PtrR-binding area at *fnrS*, which shares a common consensus with the identified DNA motif at *sad/ptrR*. We created multiple alignments of the upstream DNA sequences of closely related species with the beginning of the *E. coli* gene for *fnrS* and these binding sites corresponded to the ChIP-exo protected areas. The binding sites TTCACGAATCGaGAA, TTCtCGATTCGTGAA, and TgaAtGcAaCGTcAA were found for *ptrR, sad(yneI)*, and *fnrS*, respectively.

Experimental assessment of the computationally predicted PtrR DNA-binding site TTCtCGATTCGTGAA in the *sad* promoter region has been facilitated using a PtrR-binding fluorescent polarization (FP) assay (Fig.6B). The recombinant overproduced PtrR was obtained using a strain from the ASKA collection. The PtrR purification procedure is described in Supplemental materials. The binding of the purified refolded PtrR protein to synthetic DNA fragments containing the predicted PtrR-binding site was assessed using FP in the assay mixture containing 10 mM urea (Fig. 6B). Specific binding of PtrR to the DNA (5’-GGGTTCTCGATTCGTGAAGGG-3’) was detected at 0.6 uM of PtrR in contrast to the negative control (PhrR) ^24^. The fluorescent polarization assay showed binding for the PtrR to the predicted *sad* binding site and the addition of GABA leads to dissociation of PtrR from the fluorescently labeled DNA (Fig. 6B), suggesting a regulatory function for Ptr/GABA catabolism for energy. If the Ptr utilization pathway intermediate, GABA, accumulates, PtrR de-repress *sad* and *fnrS*. The upregulation for *sad* and *glsB* had been detected for the *yneJ* mutant strain for adapted *E. coli* MG1655 mutant (deletion *menF-entC-ubiC*) (Fig. 5D) ^23^.

An *E. coli* BW25113 (WT) and *ptrR* mutant strain growth phenotype on glutamate as the nitrogen source in minimal medium (glycerol as the carbon source) has been detected for the growth. Cells showed a growth phenotype when the *ptrR* gene was deleted under the starvation conditions (Fig. S9); a decrease in the growth rate was observed for the *ptrR* mutant. The mRNA level for *sad* was higher in the *ptrR* mutant at these conditions (Table S2, Table 4). The *ptrR* mutation led to the upregulation of 121 genes, with almost 50% of them encoding transporters and enzymes related to amino acid metabolism; including L-histidine and L-methionine biosynthetic pathway genes *hisABCDFHI* and *metABEFLR*, respectively.

The phenotype microarray using Biolog PM2A was tested under the microaerobic conditions for the *ptrR* mutant phenotype and the *E. coli* WT BW25113 strain. The phenotype using Glu as the energy/nitrogen source was minimal when D-glucosamine or dihydroxyacetone was the carbon source. The *ptrR* mutant defect in growth/respiration with glycine, L-ornithine, or gamma-hydroxybutyrate was observed using M9 medium with L-glutamate as the sole nitrogen source. Phenotypes for the *ptrR* mutant with D-tagatose, oxalomalic acid, gamma-hydroxybutyrate, glycine, and L-alaninamide were observed under the same conditions with Glu as a supplement (Fig S10). The regulatory effect of PtrR during aerobic growth with putrescine/Glu as the nitrogen source was detected for M9 medium with glycerol as the primary carbon source. The growth of BW25113 (WT) as well as *ptrR, yneH (glsB)* null mutant strains are shown in Fig. 5B-C. The *E. coli* WT strain had a longer lag-phase compared to the *ptrR* mutant. A growth deficiency for a *glsB* mutant was observed under the same conditions, suggesting a functional relationship between GlsB (YneH) and Sad, encoding genes conserved in genome clusters with *ptrR*. The effect of a *yneH* deletion was substantial as cells approached the stationary phase.

### PtrR-dependent regulation during growth with Ptr/Glu as nitrogen sources and antibiotic resistance

The *E. coli* WT and *ptrR* mutant were grown aerobically in M9 medium with 20 mM Glu and Ptr as nitrogen sources and 0.2 % glycerol. To determine the effect of the *ptrR* deletion mutation, the cells were collected at the log-phase, and total mRNA was purified (see Materials and methods). PuuR and PuuADE, SodB, and two copper related transport systems’ mRNA levels increased in the *ptrR* mutant strain (Table 3). SodB (superoxide dismutase) mRNA was increased in *ptrR* mutant and SodB produced H_2_O_2_. The *copA* and *cus* operons are regulated by the CusSR and HprRS system. H_2_O_2._is the effector for HprRS and likely has a transcriptional effect for the *copA* and *cus* system.

Antibiotic resistance induced by *ptrR* mutation in *E. coli* BW25113 was detected. We propose that PtrR negatively controls the FnrS small RNA, which is involved in regulation of MarA mRNA. MarA is a global regulator of *E. coli* genes involved in resistance to antibiotics, oxidative stress, organic solvents, and heavy metals ^25^. We tested the *ptrR* mutant and wild type *E. coli* strains for antibiotic resistance using the Biolog plate 11C ^26^. Since FnrS is under positive control of the global anaerobic regulator Fnr, *E. coli* was grown under microaerobic conditions. The *ptrR* mutant showed increased resistance to high concentrations of demeclocycline, a tetracycline group antibiotic, which survived after 42 hours, in contrast to the wild type. We also detected the increased resistance of the *ptrR* mutant to chlorotetracycline (another tetracycline analog) (Fig. S11). However, with other antibiotics tested, no significant difference in growth of the mutant and wild type strains was observed.

## DISCUSSION

In this study, we applied a systems approach to characterize the transcriptional responses of seven putative LTFs: YbdO, YbeF, YbhD, YcaN, YiaU, YgfI, and YneJ (PtrR) (Table 5). The transcriptional response for the deletion of each LTF had been detected by RNAseq in M9 minimal medium supplemented by 7 mM L-threonine for all yTFs, except PtrR. The transcriptional analysis for the *ptrR* deletion mutant was detected in the M9 medium with Ptr and/or Glu as nitrogen source. For LTFs, conserved adjacent genes had been shown to be differentially expressed. For example, gene clusters encoding *ybdNM* and citrate utilization genes *citCDEF* (citrate lyase) were detected as transcriptionally regulated in the *ybdO* deletion mutant. The YbdO DNA-binding site upstream of *ybdO* has been predicted and confirmed by ChIP-exo assay, suggesting autoregulation. YbdO has been shown to be important for the growth in minimal medium supplemented by citrate at acidic conditions, suggesting citrate lyase regulation. Additionally, flagella biosynthesis genes (FlhDC regulon) were differentially expressed in the mutant. The *ybdO* mutant in *E. coli* BW25113 fitness phenotype had been previously shown for motility in LB medium. Additionally, YbdO is important for *E. coli* BW25113 fitness in minimal medium with glycolate as the carbon source (fit.genomics.lbl.gov), and D-glycine as the nitrogen source. YbeF is conserved in the gene cluster with citrate lyase encoding genes. The *ybeF* deletion leads to *lrhA* gene downregulation and upregulation of FlhDC regulated genes and, accordingly, the FlhDC iModulon.

The YgfI (DhfA) DNA-binding site upstream of the *dhaKLM* operon for dihydroxyacetone phosphotransferase was detected by ChIP-exo. The transcriptome analysis shows that DhfA was important for regulation of *dhaKLM, pflB, hycBCDEFG, narZYWV* and *adhB*. The *dhfA* mutant growth phenotype using glycerol as carbon source had been detected (Fig.3C). The common DNA-binding motif upstream of the *dhak, pflB, adhE, hycB*, and *narZ* genes was found, but future experiments to confirm DhfA binding to the predicted DNA-binding sites are essential (Table 5). The PflB and *hycBCDEFG* encoded hydrogenlyase are involved in pyruvate and Thr utilization as energy source (Fig. 3F). The *dhfA* mutant growth deficiency on glucose as the carbon source in minimal medium at microaerobic conditions had been shown, but supplementation by Thr reduced the growth phenotype (Fig. S7).

ChIP-exo detected multiple YiaU DNA-binding sites (Fig. 4A). The RNAseq transcriptomic analysis and ChIP-Exo (YiaU DNA-binding) additionally detected direct regulation of the *waaPSBOJYZU* operon, *yjiT, gltB* and *adeP* genes. *yiaU* deletion mutant transcriptomic analysis suggested the regulation of *yiaT*, the conserved adjacent gene to *yiaU*, which is divergently transcribed (Fig. 4D).

The *ybhD* adjacent gene *ybhH* conserved in enterobacterial genomes was upregulated in the *ybhD* deletion mutant. The *ybhH, ybhI*, and *ybhD* genes are conserved adjacent genes and potentially regulated by Nac (EcoCyc). YbhI is the putative tricarboxilate transporter, homologous to 2-oxoglutarate/malate translocator, (id. 35%) (Fig. S8) ^27^. We detected the *ybhD* deletion strain growth phenotype in the M9 minimal medium with glycerol (carbon source), supplemented by L-malate (Fig. S9), but not in the absence of L-malate, suggesting that the de-repressed *ybhI* and *ybhH* genes are probably involved in L-malate utilization (Table 5).

We detected that PtrR (YneJ) is the transcriptional regulator for Sad and the small RNA, FnrS. PtrR was predicted to be a repressor of the *fnrS* gene, encoding a small regulatory RNA (Table 5). We demonstrated PtrR binding to the predicted DNA-binding site. According to RNA-Seq data, PtrR is a repressor for *sad* under the nutrient limitation-stress conditions. The *ptrR* gene deletion effect was shown by growth phenotype (aerobically) and phenotype microarray data (micro-aerobically).

PtrR-mediated regulation appears to be important for Ptr utilization as an energy source. A pleiotropic effect of the PtrR-dependent regulation of the *sad* gene under nitrogen/carbon starvation has been investigated and discussed. The known stress/starvation sigma σ^S^-controlled *csiD-ygaF*-*gabDTP* region is related to GABA utilization, while Sad is important for Ptr utilization ^20,28–30^.

Extracellular Ptr alters the OmpF porin charge and pore size, resulting in partial pore closure and a consequent decrease in outer membrane permeability ^13,31^. Our results demonstrated that PtrR is important for the growth of the *E. coli* BW25113 strain with Glu as the sole nitrogen source and glycerol as the carbon source and resistance to the tetracycline group of antibiotics (i.e., demeclocycline and chlortetracycline), but not to chloramphenicol, erythromycin, and other antibiotics. PtrR is potentially important for the regulation of the highly conserved, anaerobically induced small RNA-*fnrS*, which is likely important for regulation under anaerobic growth conditions ^32^. Interestingly, a *ptrR* mutant was shown previously to be resistant to bacteriophage lambda infection ^33^ and we found that PtrR is potentially related to tetracycline resistance ^34^. ChIP-exo and RNAseq results have been analyzed, providing a hypothesis for the physiological functions of YneJ (re-named PtrR, putrescine related regulator), YgfI (re-named DhfA, dihydroxyacetone phosphotransferase and formate utilization activator), and YbdO (re-named CtrR, citrate utilization related regulator).

Taken together, the systems analysis of the *E. coli* BW25113 and MG1655 strains’ transcriptomic data, ChIP-exo DNA-binding data for LTF, and the regulated biochemical pathway reconstruction and fitness/phenotype of the LTF deletion mutants strains produce fruitful hypotheses for the yTF function prediction that is important for TRN reconstruction in *E. coli*.

## METHODS

### RNA sequencing

The wild type BW25113 strain was grown as a control for the isogenic *ptrR* mutant strain. Pre-cultures were obtained by scraping frozen stocks and growing the cells in LB medium. Cells were washed twice with M9 medium and inoculated to an OD_600_ of 0.05. The cells were collected at an OD_600_ of 0.9 (only *ycaN, yiaU, ybhD* mutants and the WT control were collected at OD_600_ of 2.0; late-exponential phase) and were harvested using the Qiagen RNA-protect bacteria reagent according to the manufacturer’s specifications. Pelleted cells were stored at -80°C, and after cell resuspension and partial lysis, they were ruptured with a beat beater; the total RNA was extracted using a Qiagen RNA purification kit. After total RNA extraction and subsequent ribosomal RNA removal, the quality was assessed using an Aglient Bioanalyser using an RNA 6000 kit. The data processing is described in Supplemental materials.

### Protein purification

The PrtR-producing strain was grown overnight, re-inoculated into 50 mL of the fresh medium, and induced with 0.6 mM IPTG after an OD_600_ of 0.6 was reached. The cells were harvested after 4 hours and lysed, the cell pellet was resuspended in the lysis buffer. Rapid purification of recombinant proteins on Ni-nitrilotriacetic acid-agarose minicolumns was performed. The protein was refolded on a mini-column, and the buffer was changed to a buffer containing 0.1M Tris-HCl, 0.1 M NaCl, 10 mM urea.

### Fluorescent polarization assay

The purified PtrR protein and 10 nM fluorescently labeled DNA fragment (5’-gggTTCTCGATTCGTGAAggg-3’) were incubated in the assay mixture. The PtrR binding assay mixture (0.1 ml) contained Tris buffer, pH 7.5, 0.1 M NaCl, 10 mM MgSO_4_, 5 mg/ml sperm DNA and 1uM of the fluorescently labeled predicted PtrR binding DNA fragment as well as 0-0.6 mM GABA. Then the PtrR protein (0-1.5 uM) was added to the assay mixture, and it was incubated for 1 hour at 30°C.

### Targeted high-performance liquid chromatography

For organic acid and carbohydrate detection, samples were collected after 4 hours for every 30-45□Jminutes. The filtered samples were loaded onto a 1260 Infinity series (Agilent Technologies) high-performance liquid chromatography (HPLC) system with an Aminex HPX-87H column (Bio-Rad Laboratories) and a refractive index detector and HPLC was operated using ChemStation software. The HPLC was run with a single mobile phase composed of HPLC grade water buffered with 5 mM sulfuric acid (H_2_SO_4_). The flow rate was held at 0.5□JmL/minute, the sample injection volume was 10 uL, and the column temperature was maintained at 45□J°C. The identities of compounds were determined by retention time comparison to standard curves of acetate, ethanol, glucose, lactate, pyruvate, formate and succinate. The peak area integration and resulting chromatograms were generated within ChemStation and compared to that of the standard curves to determine the concentration of each compound in the samples.

## TABLES

Table 1. Differentially expressed genes revealed by RNA-Seq *ybdO* deletion mutant strain and wild type *E. coli* strains during growth in M9 medium with glucose as the primary carbon source and 7 mM L-threonine as supplement.

Table 2. Differentially expressed genes revealed by RNA-Seq *ygfI* deletion mutant strain and wild type *E. coli* strains during growth in M9 medium with glucose as the primary carbon source and 7 mM L-threonine as supplement.

Table 3 Differentially expressed genes revealed by RNA-Seq of a *ptrR* knockout and wild type *E. coli* strains during growth in M9 medium with L-glutamate and putrescine as nitrogen sources and glycerol as the primary carbon source.

Table 4 Differentially expressed *ptrR* and the adjacent genes revealed by RNA-Seq of a *ptrR* knockout and wild type *E. coli* BW25113 strains during growth in M9 medium with L-glutamate as nitrogen sources and glycerol as the primary carbon source.

Table 5 Summary of the yTFs predicted function and the yTF regulated genes evolved by transcriptomic analysis and ChIP-exo DNA-binding newly characterized in this study

## Notes

### Competing Interest Statement

The authors have declared no competing interest.

