## Supplemental Table 1, Figs S1-S10 for "A systems approach discovers the role and characteristics of seven LysR type transcription factors in *Escherichia coli*"

**Supplemental materials.**

RNA sequencing. Paired-end RNA-sequencing libraries were prepared as described (38).

Raw-sequencing reads were mapped to the reference genome (NC_000913.3) of *E. coli* K-12 MG1655 using bowtie (v1.1.2) with the following options “-X 1000 -n 2 -3 3”. Transcript abundance was quantified using *summarizeOverlaps* from the R *GenomicAlignments* package (v1.18.0) with the following options: “mode = “IntersectionStrict”, singleEnd = FALSE, ignore.strand = FALSE, preprocess.reads = invertStrand”. To ensure the quality of the compendium, genes shorter than 100 nucleotides and genes with under 10 fragments per million-mapped reads across all samples were removed before further analysis. Transcripts per million (TPM) were calculated by *DESeq2* (v1.22.1). The final expression values were log-transformed log^2^(TPM + 1), referred to as log-TPM.

**ChIP-Exo experiments.** Chip-exo experiments were performed following the procedures previously described. To identify PtrR binding sites for each strain, the DNA bound to PtrR from formaldehyde cross-linked cells, collected after growth in M9, was isolated by chromatin immunoprecipitation (ChIP) with the antibodies that specifically recognize the myc tag (9E10, Santa Cruz Biotechnology), and Dynabeads Pan Mouse IgG magnetic beads (Invitrogen) were added, followed by stringent washings as described previously ^2^. ChIP materials (chromatin-beads) were used to perform on-bead enzymatic reactions of the ChIP-exo method. Briefly, the sheared DNA of the chromatin-beads was repaired by the NEBNext End Repair Module (New England Biolabs), followed by the addition of a single dA overhang and ligation of the first adaptor (5’-phosphorylated) using a dA-Tailing Module (New England Biolabs) and NEBNext Quick Ligation Module (New England Biolabs), respectively. Nick repair was performed by using the PreCR Repair Mix (New England Biolabs). Lambda exonuclease- and RecJf exonuclease-treated chromatin was eluted from the beads, and overnight incubation at 65 degrees reversed the protein-DNA cross-links. RNA- and protein-free DNA samples were used to perform primer extension and second adaptor ligation with the following modifications. The DNA samples, incubated for primer extension as described previously (38), were treated with the dA-Tailing Module and NEBNext Quick Ligation Module (New England Biolabs) for second adaptor ligation. The DNA sample, purified using the GeneRead Size Selection Kit (Qiagen), was enriched by polymerase chain reaction (PCR) using Phusion High-Fidelity DNA Polymerase (New England Biolabs). The amplified DNA samples were purified again with a GeneRead Size Selection Kit (Qiagen) and quantified using Qubit dsDNA HS Assay Kit (Life Technologies). The quality of the DNA sample was checked by running the Agilent High Sensitivity DNA Kit using an Agilent 2100 Bioanalyzer before sequencing using HiSeq 2500 (Illumina) following the manufacturer’s instructions. Each modified step was also performed following the manufacturer’s instructions. ChIP-exo experiments were performed in duplicate.

**FIGURES**

**Fig. S1** Alignment of the LTTR transcription factors in Escherichia coli K-12 MG1655. Sequence profiles of the proteins were aligned in Clustal Omega. One of the most common regulator families in prokaryotes is lysR-type transcriptional regulators (LTTR). The lysR-type HTH domain is a DNA-binding, winged-turn-helix (wHTH) domain of about 60 residues present in the N-terminal part. LTTRs are present in diverse bacterial genera, archaea and algal chloroplasts. All LTTRs contain the DNA-binding lysR-type HTH domain.

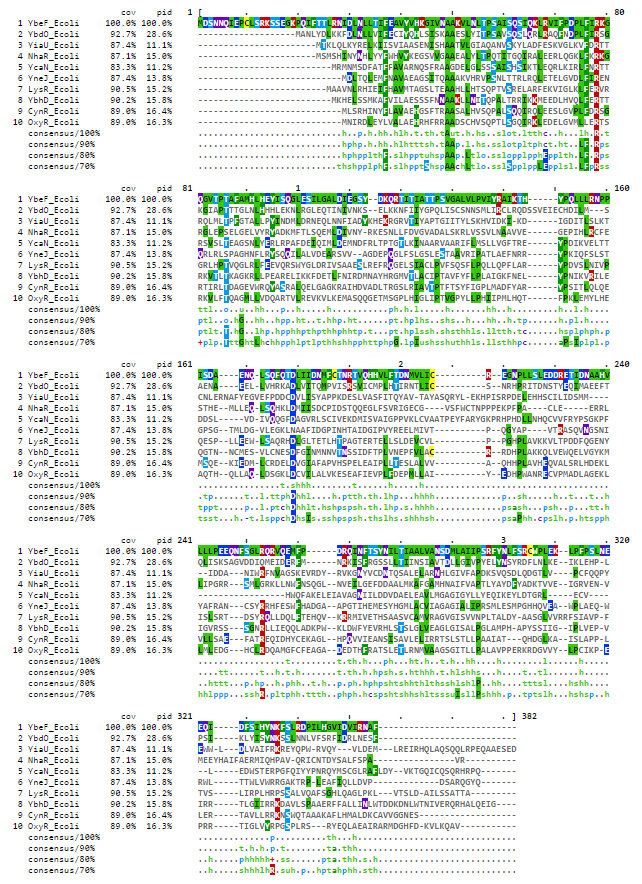

**Fig. S2** The predicted YbdO binding site (yellow backgrond) upstream of *ybdO* gene, suggesting that *ybdO* is autoregulator as supported by ChiP-exo DNA-binding and RNA-seq data (Table 2), the ybdO start of the gene is marked by red font.

display  items per page

| T | displaying 1 - 10 of 11 | next»  last» |
| --- | --- | --- |

| **ID** | **Organism** | **Annotation** |
| --- | --- | --- |
| fig\|83333.1.peg.604 | Escherichia coli K12 | LysR-family transcriptional regulator YbdO |
| fig\|1068608.3.peg.3106 | Escherichia coli XH140A | LysR-family transcriptional regulator YbdO |
| fig\|457401.3.peg.1903 | Escherichia sp. 4_1_40B | Putative LysR-family transcriptional regulator YbdO |
| fig\|556266.3.peg.3380 | Shigella sp. D9 | Putative LysR-family transcriptional regulator YbdO |
| fig\|766158.3.peg.679 | Shigella flexneri 2850-71 | Putative LysR-family transcriptional regulator YbdO |
| fig\|216599.1.peg.581 | Shigella sonnei 53G | Putative LysR-family transcriptional regulator YbdO |
| fig\|621.8.peg.5123 | Shigella boydii ATCC 9905 | Putative LysR-family transcriptional regulator YbdO |
| fig\|358708.5.peg.2095 | Shigella dysenteriae 1012 | Putative LysR-family transcriptional regulator YbdO |
| fig\|754333.3.peg.2625 | Escherichia albertii TW11588 | Putative LysR-family transcriptional regulator YbdO |
| fig\|585054.5.peg.2409 | Escherichia fergusonii ATCC 35469 | Putative LysR-family transcriptional regulator YbdO |

Bottom of Form

fig|83333.1.peg.604	Escherichia coli K12

fig|1110693.3.peg.466	Escherichia coli str. K-12 substr. MDS42

fig|457401.3.peg.1903	Escherichia sp. 4_1_40B

fig|556266.3.peg.3380	Shigella sp. D9

fig|624.753.peg.3857	Shigella sonnei strain NLAE-zl-G496

fig|373384.11.peg.637	Shigella flexneri 5 str. 8401

fig|621.221.peg.5428	Shigella boydii strain ATCC 35964

fig|564.14.peg.4142	Escherichia fergusonii strain FDAARGOS_170

fig|622.435.peg.3171	Shigella dysenteriae strain ATCC 12039

fig|208962.68.peg.595	Escherichia albertii strain 1551-2

fig|1499973.6.peg.1976	Escherichia marmotae strain HT073016

fig|67829.4.peg.1024	Citrobacter murliniae strain P080C CL

fig|67827.10.peg.3160	Citrobacter werkmanii strain FDAARGOS_364

fig|1006003.3.peg.132	Citrobacter freundii ATCC 8090 = MTCC 1658

fig|545.11.peg.827	Citrobacter koseri strain FDAARGOS_86

fig|133448.4.peg.3409	Citrobacter youngae strain L6

fig|469595.3.peg.1261	Citrobacter sp. 30_2

fig|1218086.3.peg.1181	Citrobacter sedlakii NBRC 105722

fig|57706.65.peg.4328	Citrobacter braakii strain MiY-A

fig|1218085.3.peg.3193	Citrobacter rodentium NBRC 105723

fig|1064551.5.peg.662	Salmonella enterica subsp. enterica serovar Thompson str. RM6836

```
fig|208962.68.peg.595  t-agaaaaaCTTAAcgataaaTTAAGtaaagaa-a-----------ataa
fig|1499973.6.peg.1976 t-agaaaaaCTTAAtcgtgaaTTAAGtaaaata-acataatatataaaaa
fig|564.14.peg.4142    t-agaaaatCTTAAtccaaagTTAAGcagaaaa-acataat----acata
fig|622.435.peg.3171   t-agaaaaaCTTAAtcctgagTTAAGtaaaaaa-acataat----ataaa
fig|373384.11.peg.637  t-agaaaaaCTTAAtcctgagTTAAGcaaaaaa-acataatcaataaaaa
fig|621.221.peg.5428   t-agaaaaaCTTAAtcctgagTTAAGcaaaaaa-acataatcaataaaaa
fig|556266.3.peg.3380  t-agaaaaaCTTAAtcctgagTTAAGcaaaaaa-atataatcaataaaaa
fig|624.753.peg.3857   t-agaaaaaCTTAAtcctgagTTAAGcaaaaaa-atataatcaataaaaa
fig|83333.1.peg.604    t-agaaaaaCTTAAtcctgagTTAAGcaaaaaa-acataatcaataaaaa
fig|1110693.3.peg.466  t-agaaaaaCTTAAtcctgagTTAAGcaaaaaa-acataatcaataaaaa
fig|457401.3.peg.1903  t-agaaaaaCTTAAtcctgagTTAAGcaaaaaa--cataatcaataaaaa
fig|67829.4.peg.1024   t-agttttagTTAActgttcaTTAAGatttaaatggatatgtgctaagaa
fig|67827.10.peg.3160  t-agtttaagTTAActattcaTTAAGcgtataaaagattcttaaaagaaa
fig|1006003.3.peg.132  t-agctaaagTTAAttattcaTTAAGcacacaa-acataaatttcaagaa
fig|57706.65.peg.4328  ttagtttaaaTTAAtaattcaTTAAGcacacaa-acataaatcacaagaa
fig|133448.4.peg.3409  t-agtttaagTTAAttattcaTTAAGc-cataa-acataaactctaagaa
fig|469595.3.peg.1261  t-agtttaagTTAAttattcaTTAAGc-cataa-acataaactctaagaa
fig|545.11.peg.827     t-agtattatTgAAagagaatTTAAGcttagcaaacatcactattgtaaa
fig|1218086.3.peg.1181 t-agccataCTaAAtgtcactTTAAGtttg-cattgaaaaattttattca
fig|1218085.3.peg.3193 t-agccataCTaAAactaactTTAAGtcaggcattgatttctttagctta
fig|1064551.5.peg.662  t-agtttaaaTaAActctacaTTAAGtttaataaacataactctggtaaa
                       * **      * **       *****      *                *

fig|1499973.6.peg.1976 tacataaagcca-tatcattgacttgatgaata--ttatgagaacaagca
fig|564.14.peg.4142    taaatacaacaa-cattattgaacaagtgaatgatttatggaagagacaa
fig|622.435.peg.3171   taca-acagcag-cattattgaacaaatgaatgatttatagaagagataa
fig|373384.11.peg.637  tacatacgcaaa-aaacattgattaagtgaatataccatggaagaaa-aa
fig|621.221.peg.5428   tacatacgcaaa-aaacattgattaagtgaatataccatggaagaaa-aa
fig|556266.3.peg.3380  tatatacgcaaa-aaacattgattaagtgaatataccatggaagaaa-aa
fig|624.753.peg.3857   tatatacgcaaa-aaacattgattaagtgaatataccatggaagaaa-aa
fig|83333.1.peg.604    tatatacgcaaa-aaacattgattaagtgaatatatcatggaagaaa-aa
fig|1110693.3.peg.466  tatatacgcaaa-aaacattgattaagtgaatatatcatggaagaaa-aa
fig|457401.3.peg.1903  tatatacgcaaa-aaacattgattaagtgaatatatcatggaagaaa-aa
fig|67829.4.peg.1024   tgattaataac--ttagaca--taatgaaacggaatatttcagagcaaag
fig|67827.10.peg.3160  taatttgcaat--tcaaata--tataaaaatgaaatatttccacataaaa
fig|1006003.3.peg.132  taactaccatt--taaacta--ttccacaattgaatttttcactacatta
fig|57706.65.peg.4328  tattcacaact--taaaata--tttcagaattaaatttttcaaaacatga
fig|133448.4.peg.3409  tgattagaatt--taaatta--ttcaacaattgaattatcctatacatta
fig|469595.3.peg.1261  tgattagaatt--taaatta--ttcaacaattgaattatcctatacatta
fig|545.11.peg.827     gcattggatgtg-ttgcataactataacaaatatagttatgacaatagtc
fig|208962.68.peg.595  tacataaagcca-tactattgacttgatgaata--ttatgataa-aagca
fig|1218086.3.peg.1181 gtgacaacacctaaaacattcatgcaacagataatttcatcttagaaa-t
fig|1218085.3.peg.3193 a-gatgacacggaaagcattaatgatacggatgatttcatatcaaaaagt
fig|1064551.5.peg.662  ttattgagatt--aagtcataat--catggaagataccatgataatgc--

fig|1499973.6.peg.1976 t-----atga-atggagtgcatcATGGCTAATCTTTACGACTTAAAAAAA
fig|564.14.peg.4142    t-----acaa--cggagtatgttATGGCTAATCTTTATGATCTAAAAAAA
fig|622.435.peg.3171   t-----atga--cggagtatgttATGGCGAATCTTTATGGCCTAAAAAAA
fig|373384.11.peg.637  t-----ataaaccggagtagtgtATGGCCAATCTCTACGACTTGAAAAAG
fig|621.221.peg.5428   t-----ataaaccggagtagtgtATGGCCAATCTCTACGACTTGAAAAAG
fig|556266.3.peg.3380  t-----ataa-ccggagtagtgtATGGCCAATCTCTACGACTTGAAAAAG
fig|624.753.peg.3857   t-----ataa-ccggagtagtgtATGGCCAATCTCTACGACTTGAAAAAG
fig|83333.1.peg.604    t-----ataa-ccggagtagtgtATGGCCAATCTCTACGACTTGAAAAAG
fig|1110693.3.peg.466  t-----ataa-ccggagtagtgtATGGCCAATCTCTACGACTTGAAAAAG
fig|457401.3.peg.1903  t-----ataa-ccggagtagtgtATGGCCAATCTCTACGACTTGAAAAAG
fig|67829.4.peg.1024   c-taagacaggaatggata---tATGGCTAACCTTTACGACCTCAAAAAA
fig|67827.10.peg.3160  t-aacaatgagaatataca---tATGGCTAACCTCTATGACCTTAAAAAA
fig|1006003.3.peg.132  c-catcacaggaaggtacg---cATGGCTAACCTCTATGACCTCAAAAAG
fig|57706.65.peg.4328  c-catcacaggaaggtatg---tATGGCTAACCTCTATGACCTTAAAAAG
fig|133448.4.peg.3409  c-catcacaggaaggtatg---cATGGCTAACCTCTATGACCTCAAAAAA
fig|469595.3.peg.1261  c-catcacaggaaggtatg---cATGGCTAACCTCTATGACCTCAAAAAA
fig|545.11.peg.827     a-caaaacaggaagaaaaaaa-cATGGCTAATCTCTACGATCTCAAAAAA
fig|208962.68.peg.595  tgaatcatga-atggagtgcgtcATGGCTAATCTGTATGATTTAAAGAAA
fig|1218086.3.peg.1181 g------gaaacaggaaaaaaacATGGCTAACCTTTACGATCTCAAAAAA
fig|1218085.3.peg.3193 g------agaaaaggaaaaacacATGGCTAACCTTTACGATCTCAAAAAG
fig|1064551.5.peg.662  -----------cgggaacatactATGGCTAACCTTTACGACCTTAAAAAA
                                              ***** ** ** ** *   * ** **
```

**Fig S3** YbeF and ybdO gene clustering with citCDEFXGT (citrate lyase) and citAB(transcriptional regulator)in E. coli, Salmonella, Citrobacter sp.

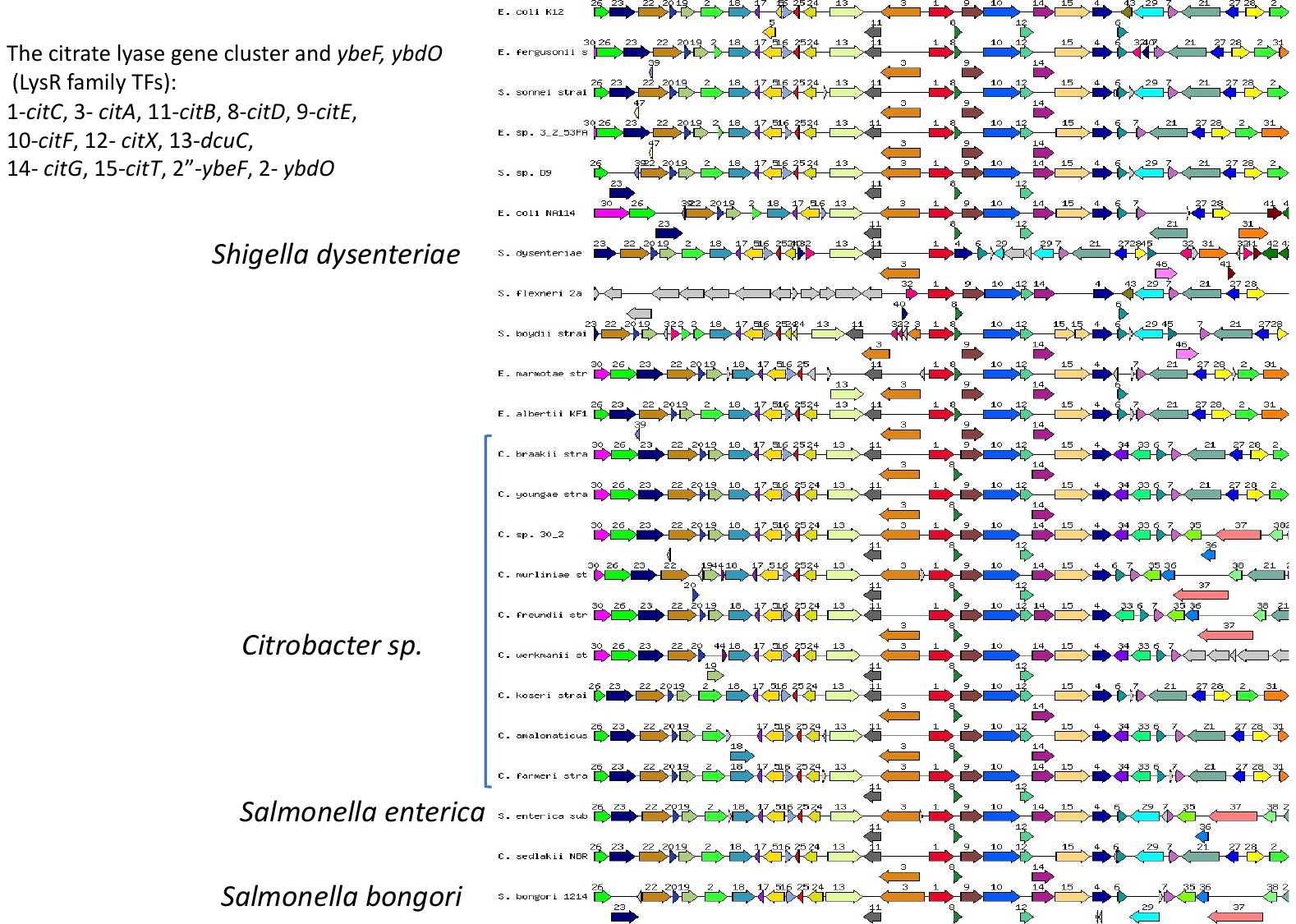

FlhDC

|  |  |
| --- | --- |
| WT | 0 |
| ybdO | 6.86436 |
| ybeF | 12.2195 |
| ygfI | 14.1658 |
| WT_LL | 1.20281 |
| ycaN_LL | 14.1658 |
| yiaU_LL | 7.20427 |

**Fig. S4** FlhDC regulon and additional flagella related genes form “FlhDC” iModulon (iModulonDB), transcriptional activity for yTF mutant strains in mid-exponential (WT, ybdO, ybeF, ygfI and late exponential (WT_LL, ycaN_LL, yiaU_LL) growth phase.

**Fig. S5A** *YbhJ* (red) genome context with the *ybhD* (dark purple), *ybhI* (dark blue), Gnl-2-oxoglutarate/malate translocator precursor and *ybhH* (dark salad green).

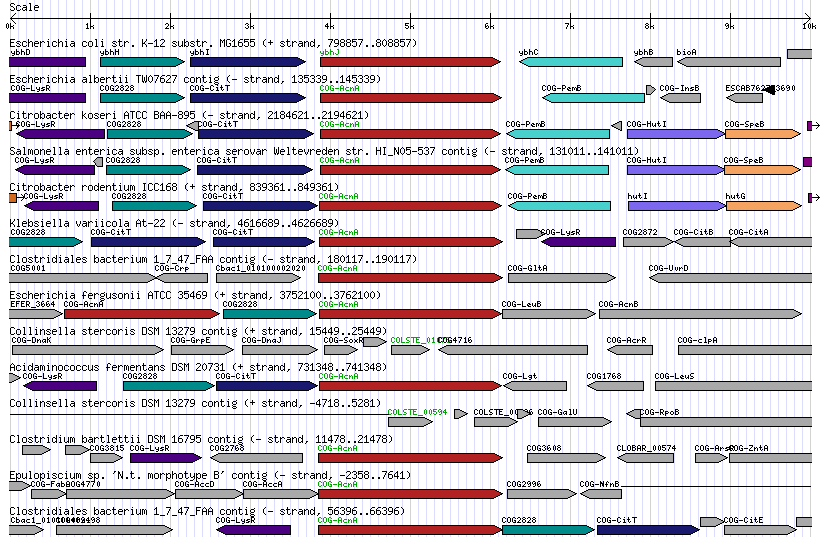

**Fig S5B YbhI alignment with the Gnl-2-oxoglutarate/malate translocator precursor from Spinach (TCDB).**

>gnl|BL_ORD_ID|13007 gnl|TC-DB|Q41364|2.A.47.3.1 2-OXOGLUTARATE/MALATE

TRANSLOCATOR PRECURSOR - Spinacia oleracea (Spinach). Length = 569

Score = 810, Bits(316), Expect = 9.11157e-101

Identities=35.7%, Positives=56.4%

Query:	5SLWKLILILAIPCIIGFMPAPAGLSELAWVLFGIYLAAIVGLVIKPFPEPVVLLIAVAAS	64

S+	L+ +	I I	F+ P P	G+S	AW L	I+ L+	IVG++	+P	P	V	L+	+ AS

```
Sbjct:103SIKPLLASILTGVIIWFIPTPEGVSRNAWQLLAIFLSTIVGIITQPLPLGAVALMGLGAS162

Query:	65 MVVVGNLSDGAFKTTAVLSGYSSGTTWLVFSAFTLSAAFVTTGLGKRIAYLLIGKIGNTT 124
```

++	AF	S +	WL+	AF +	F+	TGLG	RIAY +	G+++

```
Sbjct: 163 VLTKTLTFSAAF------SAFGDPIPWLIALAFFFARGFIKTGLGNRIAYQFVKLFGSSS 216

Query:	125 LGLGYVTVFLDLVLAPATPSNTARAGGIVLPIINSVAVALGSE-PEKSPRRVGHYLMMSI 183
```

LGLGY	VF + +. LAPA	PS +. ARAGGI	LP++	S+ +. A	GS	+ +	R++G +. LM++

```
Sbjct:	217 LGLGYSLVFSEALLAPAIPSVSARAGGIFLPLVKSLCIACGSNVGDGTERKLGAWLMLTC 276

Query:	184 YMVTKTTSYMFFTAMAGNILALKMINDILHLQISWGGWALAAGLPGIIMLLVTPLVIYTM 243
```

+ + +. S	MF TAMA N	L+ + + +	I	W	WA AA +. PG++ L+ V	PL++ Y +

```
Sbjct:	277 FQTSVISSSMFLTAMAANPLSATLTFNTIGKAIGWMDWAKAAFVPGLVSLIVVPLLLYVV 336

Query:	244 YPPEIK-KVDNKTIAKAGLAELGPMKIREKMLLGVFVLALLGWIFSKSLGVDESTVAIVV 302
```

YPPEIK	D	+	AK	L ++. GPM	E ++	+. L +	W+F	LGVD	T AI+

```
Sbjct:	337 YPPEIKSSPDAPRLAKEKLDKMGPMTKNESIMAVTLLLTVGLWVFGGKLGVDAVTAAILG 396

Query:	303 MATMLLLGIVTWEDVVKNKGGWNTLIWYGGIIGLSSLLSKVKFFEWLAE-VFKNNLAFDG 361
```

++	+L+	G+VTW++ +	W+TL W+ +I ++	L+K	W +E	V	K

```
Sbjct:	397 LSVLLITGVVTWKECLAESVAWDTLTWFAALIAMAGYLNKYGLITWFSENVVKVVGGLGL 456

Query:	362 HGNVAFFVIIFLSIIVRYFFASGSAYIVAMLPVFAMLANVSGAPLMLTALALLFSNSYGG 421
```

++F	V++ L	YFFASG+A+I	AM	F +A+	G	P	L	A+	L	F ++	G

```
Sbjct:	457 SWQMSFGVLVLLYFYSHYFFASGAAHIGAMFTAFLSVASALGTPPFLAAIVLSFLSNLMG 516

Query:	422 MVTHYGGAAGPVIFGVGYNDIKSWWLVGAVLTILTFLVHITLGVWWWNMLIGW	474
```

+THYG +	PV. +G	Y +	WW	G +++I+	++ + +G	WW +	W

Sbjct:	517 GLTHYGIGSAPVFYGANYVPLPQWWGYGFLISIVNLIIWLGVGGLWWKAIGLW	569

**Fig. S6 The** growth measurement of the *ybhD* mutant  (orange and grey line) compared to the wild type BW25113 strain of *E. coli* (blue and yellow line). Growth was measured in 96-well plates with glucose or glycerol as the carbon source (in the presence and the absence of 10 mM malate).

M9, 0.4 % glycerol, L-malate 10mM

|  | WT-Malate | ybhD-Malate |
| --- | --- | --- |
| 0 | 0.142 | 0.139 |
| 0.0138889 | 0.153 | 0.147333 |
| 0.0277778 | 0.1665 | 0.158 |
| 0.0416667 | 0.1785 | 0.168333 |
| 0.0555556 | 0.1865 | 0.178 |
| 0.0694444 | 0.199 | 0.186 |
| 0.0833333 | 0.2185 | 0.199667 |
| 0.0972222 | 0.244 | 0.22 |
| 0.111111 | 0.2725 | 0.247 |
| 0.125 | 0.3045 | 0.279667 |
| 0.138889 | 0.338 | 0.311333 |
| 0.152778 | 0.3715 | 0.313667 |
| 0.166667 | 0.4115 | 0.331667 |
| 0.180556 | 0.4355 | 0.368333 |
| 0.194444 | 0.4595 | 0.420333 |
| 0.208333 | 0.4865 | 0.479 |
| 0.222222 | 0.51 | 0.546667 |
| 0.236111 | 0.536 | 0.624 |
| 0.25 | 0.58 | 0.694667 |
| 0.263889 | 0.6055 | 0.751667 |
| 0.277778 | 0.6335 | 0.806667 |
| 0.291667 | 0.664 | 0.858 |
| 0.305556 | 0.69 | 0.904333 |
| 0.319444 | 0.717 | 0.946667 |
| 0.333333 | 0.746 | 0.983 |
| 0.347222 | 0.769 | 1.01533 |
| 0.361111 | 0.7955 | 1.04333 |
| 0.375 | 0.8175 | 1.08467 |
| 0.388889 | 0.841 | 1.10667 |
| 0.402778 | 0.8585 | 1.12233 |
| 0.416667 | 0.882 | 1.14067 |
| 0.430556 | 0.8955 | 1.16167 |
| 0.444444 | 0.9165 | 1.17667 |
| 0.458333 | 0.9325 | 1.18533 |
| 0.472222 | 0.935 | 1.192 |
| 0.486111 | 0.925 | 1.212 |
| 0.5 | 0.917 | 1.225 |
| 0.513889 | 0.91 | 1.22233 |
| 0.527778 | 0.904 | 1.21967 |
| 0.541667 | 0.8985 | 1.21667 |
| 0.555556 | 0.894 | 1.21433 |
| 0.569444 | 0.8905 | 1.21167 |
| 0.583333 | 0.887 | 1.209 |
| 0.597222 | 0.885 | 1.20633 |
| 0.611111 | 0.883 | 1.20333 |
| 0.625 | 0.8815 | 1.19933 |
| 0.638889 | 0.8805 | 1.19667 |
| 0.652778 | 0.8795 | 1.19333 |
| 0.666667 | 0.8795 | 1.19033 |
| 0.680556 | 0.8775 | 1.18667 |
| 0.694444 | 0.878 | 1.18367 |
| 0.708333 | 0.8775 | 1.18067 |

M9 medium, 0.4% glycerol carbon source, L-malate 10 mM

M9 glucose, glycerol

|  | WT | ybhD | WT | ybhD |
| --- | --- | --- | --- | --- |
| 0 | 0.141333 | 0.136333 | 0.150333 | 0.145333 |
| 0.0138889 | 0.145 | 0.139667 | 0.169 | 0.162 |
| 0.0277778 | 0.149667 | 0.143333 | 0.185333 | 0.179 |
| 0.0416667 | 0.153667 | 0.147 | 0.200333 | 0.194 |
| 0.0555556 | 0.158 | 0.151 | 0.219667 | 0.211667 |
| 0.0694444 | 0.162 | 0.154667 | 0.248 | 0.239333 |
| 0.0833333 | 0.166333 | 0.158333 | 0.284 | 0.278333 |
| 0.0972222 | 0.170667 | 0.161667 | 0.326667 | 0.326667 |
| 0.111111 | 0.175 | 0.165667 | 0.370667 | 0.380667 |
| 0.125 | 0.179333 | 0.169 | 0.411333 | 0.423667 |
| 0.138889 | 0.183667 | 0.172667 | 0.474333 | 0.438667 |
| 0.152778 | 0.189 | 0.176667 | 0.523 | 0.500333 |
| 0.166667 | 0.194667 | 0.181667 | 0.598 | 0.578 |
| 0.180556 | 0.198333 | 0.185 | 0.679333 | 0.662333 |
| 0.194444 | 0.204333 | 0.189 | 0.753333 | 0.738333 |
| 0.208333 | 0.212 | 0.194 | 0.820333 | 0.804333 |
| 0.222222 | 0.221667 | 0.2 | 0.883667 | 0.87 |
| 0.236111 | 0.233 | 0.208 | 0.941 | 0.93 |
| 0.25 | 0.247333 | 0.217333 | 1.01333 | 0.974333 |
| 0.263889 | 0.262 | 0.227 | 1.019 | 1.039 |
| 0.277778 | 0.28 | 0.238333 | 1.03133 | 1.03667 |
| 0.291667 | 0.300667 | 0.253 | 1.036 | 1.04567 |
| 0.305556 | 0.326 | 0.269333 | 1.03867 | 1.04333 |
| 0.319444 | 0.352333 | 0.288 | 1.03167 | 1.044 |
| 0.333333 | 0.382333 | 0.310667 | 1.027 | 1.042 |
| 0.347222 | 0.414667 | 0.336333 | 1.02067 | 1.033 |
| 0.361111 | 0.451667 | 0.364667 | 1.01567 | 1.02767 |
| 0.375 | 0.49 | 0.396667 | 1.012 | 1.023 |
| 0.388889 | 0.531333 | 0.432333 | 1.01033 | 1.021 |
| 0.402778 | 0.574333 | 0.471667 | 1.01033 | 1.02033 |
| 0.416667 | 0.618333 | 0.516333 | 1.01467 | 1.02133 |
| 0.430556 | 0.661333 | 0.562667 | 1.02067 | 1.02167 |
| 0.444444 | 0.696 | 0.613 | 1.02667 | 1.02133 |
| 0.458333 | 0.729333 | 0.659667 | 1.03233 | 1.024 |
| 0.472222 | 0.763667 | 0.702 | 1.038 | 1.02867 |
| 0.486111 | 0.792667 | 0.741 | 1.04267 | 1.032 |
| 0.5 | 0.816333 | 0.776333 | 1.04833 | 1.03433 |
| 0.513889 | 0.832667 | 0.808 | 1.053 | 1.03833 |
| 0.527778 | 0.844 | 0.837333 | 1.05633 | 1.04233 |
| 0.541667 | 0.849667 | 0.863667 | 1.06133 | 1.047 |
| 0.555556 | 0.852667 | 0.888333 | 1.064 | 1.05167 |

|  | ygfI_1 | WT_2 |
| --- | --- | --- |
| 0 | 0.001 | 0.001 |
| 3 | 0.042 | 0.09 |
| 4 | 0.06 | 0.12 |
| 5 | 0.066 | 0.45 |
| 6 | 0.16 | 0.54 |
| 7 | 0.22 | 0.64 |

M9 glucose, Thr 7mM

|  | ygfI_1 | ygfI_2 | BW25113 WT_1 | BW25113 WT_2 |
| --- | --- | --- | --- | --- |
| 0 | 0.1 | 0.1 | 0.1 | 0.1 |
| 2 | 0.18 | 0.18 | 0.19 | 0.19 |
| 4.5 | 0.23 | 0.25 | 0.45 | 0.47 |
| 5 | 0.35 | 0.38 | 0.63 | 0.65 |
| 6 | 0.51 | 0.55 | 0.8 | 0.81 |
| 8.5 | 1 | 1 | 1.3 | 1.35 |

**Fig. S7** The microaerobic growth for the ygfI deletion mutant (blue,orange) and E. coli BW25113 (red, yellow) in M9 glucose medium (A) and the same medium supplemented with 7 mM L-threonine.

**Fig. S8** The predicted YneJ DNA-binding site (red font) upstream of *yneJ* gene, ChIP-Exo detected DNA is marked with green background**.**

```
                                  yneI<-
fig|457401.3.peg.516   CCGGAGTAATGGTCATcgggg----tatctcctttatgagtcatggtatgaagatacgcagatttactct
fig|1050617.5.peg.1967 CCGGAGTAATGGTCATcgggg----tatctcctttatgagtcatggtatgaagatacgcagatttactct
fig|83333.1.peg.1513   CCGGAGTAATGGTCATcgggg----tatctcctttatgagtcatggtatgaagatacgcagatttactct
fig|556266.3.peg.269   CCGGAGTAATGGTCATcgggg----tatctcctttatgagtcatggtatgaagatacgcagatttactct
fig|344609.11.peg.1857 CCGGAGTAATGGTCATcgggg----tatctcctttatgagtcatggtatgaagatacgcagatttactct
fig|216599.12.peg.1800 CCGGAGTAATGGTCATcgggg----tatctcctttatgagtcatggtatgaagatacgcagatttactct
fig|300267.13.peg.1966 CCGGAGTAATGGTCATcgggg----tatctcctttatgagtcatggtatgaagatacgcagatttactct
fig|564.14.peg.3012    CCGGAGTAATGGTCATcgggg----tatctcctttatgagtcatggtatgaagatacgcagatttactct
fig|1499973.6.peg.993  CCGGAGTGATGGTCATcgggg----tatctccttcatgagtcagggtaaaaagatacgcagatttactct
fig|208962.36.peg.1540 CCGGAGTGATGGACATcagga----tatctccattgtgagtgatatctccaggatacgcagatttactcc
fig|1006003.3.peg.577  CTGCTGTGATAGTCATtaagtgc-ctctctttatcgcagggtatggttatatcatgagcatatttactct
fig|1173691.3.peg.4273 CTGCTGTGATAGTCATtaagtgc-ctctctttatcgcagggtatggttatatcatgagcatatttactct
fig|67829.4.peg.3175   CTGCTGTGATGGTCATaggtcttactctctttatcgcaggatatggttatatcatgagcgtatttactct
fig|545.23.peg.4274    CAGGTGTCATCGTCATcggtttgtctctctacgttgcggga-atgcctgcataatgggcgcatttactct
fig|35703.30.peg.4088  CCGGTGAAATCGTCATacag-----tctctccgtcgttcttgatgtcatcatcatggacgcgtttactct
                       * *  *  ** * ***         * ***   *        *       *  **   *   *******
			   YneJ binding area     YneJ binding site   -> yneJ
fig|457401.3.peg.516 tgctttaaaatgaataatattaagccacttattcacgaatcgagaatgctATGGATCTGACCCAACTGGAGATGTT
fig|1050617.5.peg1967tgctttaaaatgaataatattaagccacttattcacgaatcgagaatgctATGGATCTGACCCAACTGGAGATGTT
fig|83333.1.peg.1513 tgctttaaaatgaataatattaagccacttattcacgaatcgagaatgctATGGATCTGACCCAACTGGAGATGTT
fig|556266.3.peg.269 tgctttaaaatgaataatattaagccacttattcacgaatcgagaatattATGGATCTGACCCAACTGGAGATGTT
fig|344609.11peg.1857tgctttaaaatgaataatattaagccacttattcacgaatcgagaatattATGGATCTGACCCAACTGGAGATGTT
fig|216599.12peg.1800tgctttaaaatgaataatattaagccacttattcacgaatcgagaatgttATGGATCTGACCCAACTGGAGATGTT
fig|300267.13.peg1966tgctttaaaatgaataatattaagccacttattcacaaatcgagaatattATGGATCTGACCCAACTGGAGATGTT
fig|564.14.peg.3012  tgctttaaaatgaataatattaagccacttattcacgaatcgagaatgttATGGATCTGACCCAACTGGAGATGTT
fig|1499973.6.peg.993tactttaaaatgaataatattaagccacttattcacgaattgagaatactATGGATCTAACCCAACTGGAGATGTT
fig|208962.36.peg1540ttcgtaaaaatgaataatattaagtcacttattcacaaatcgagaatgccATGGACCTGACCCAGTTAGAGATGTT
fig|1006003.3.peg.577ttctgaaaaatgaataatattaaccataccattcacgaatagagaaaattATGGATCTGACGCAGCTGGAAATGTT
fig|1173691.3.peg4273ttctgaaaaatgaataatattaaccataccattcacgaatagagaaaattATGGATCTGACGCAGCTGGAAATGTT
fig|67829.4.peg.3175 ttatgaaaaatgaataatattaaccacaccattcacgaatagagaaaacaATGGATCTGACGCAACTGGAAATGTT
fig|545.23.peg.4274  ttctgaaaaatgaataatattaagcactgcgttcacaaaaagagaaaatcATGGATCTGACTCAGCTGGAAATGTT
fig|35703.30.peg.4088ttccgaaaaatgaataatattaaccaaaccattcacgaagagagaaaaccATGGATCTGACCCAACTGGAGATGTT
                     *     *****************        ***** **  *****      **** ** ** **  * ** ****
```

**2^nd^ branch of YneJ in Serratia / Yersinia / Erwinia**

```
fig|1206776.4.peg.1570catgccgcagtattaaa-ctcacaaaaaatgaataatactcatcaagttcatcacgaaaagagaagac-cATGGA
fig|399741.3.peg.1566 catgccgcagtattaaa-ctcacaaaaactgaataatactcaccaagttcatcacgaaaagagaagac-aATGGA
fig|615.1.peg.4875    catgccgcagtattaaa-ctcacaaaaactgaataatagttatcaagttcatcacgaaacgagaatat-tATGGA
fig|349967.3.peg.2685 catgcctgagcattaga-atcacaaaaaatgaataatagtcactaagttcatcacaaaaagagaacacttATGGA
fig|349968.3.peg.1236 catgcctgagcattaga-atcacaaaaaatgaataatactcattaagttcatcacaaaaagagaatacatATGGA
fig|349966.3.peg.2345 catgcctgagcattaga-atcacaaaaaatgaataattctcactaagttcatcacaaaaagagaacgattATGGA
fig|349965.3.peg.1395 catgcctgagcattaga-atcacaaaaaatgaataatgctcattaagttcatcacaaaaagagaacacctATGGA
fig|630.2.peg.2749    catgcctgagcattaga-atcacaaaaaatgaataatactcactaagttcatcacggaaagagaATGCTTATGGA
fig|187410.1.peg.2620 catgcctgagtattaga-atcacaaaaaatgaataatagtcattaagttcatcacaaaaagagaacatctATGGA
fig|273123.1.peg.1629 catgcctgagtattaga-atcacaaaaaatgaataatagtcattaagttcatcacaaaaagagaacatctATGGA
fig|218491.3.peg.2039 cacagcttgccgctacacttcacaaaaaatgaataatcaaacaccagttcattacgaaaagagaaaac-cATGGA
                      **   *       ** *  ********* ********        ******* **  ** ****      *****
				    -> yneJ
```

**Fig.S9 A.** The aerobic growth for the ptrR (yneJ) (dark blue line) deletion mutant and *E. coli* BW25113 (orange line) in modified M9 medium glycerol (0.1%) as carbon source and 20 mM L-Glutamate (nitrogen source), and in 96-well plates and the same medium (with glycerol 0.4%) in culture tubes.

**B.** The phenotype in plates PM2 for the ptrR (yneJ) deletion mutant and *E. coli* BW25113 in M9 medium without carbon source, supplemented by L-glutamate (20mM)

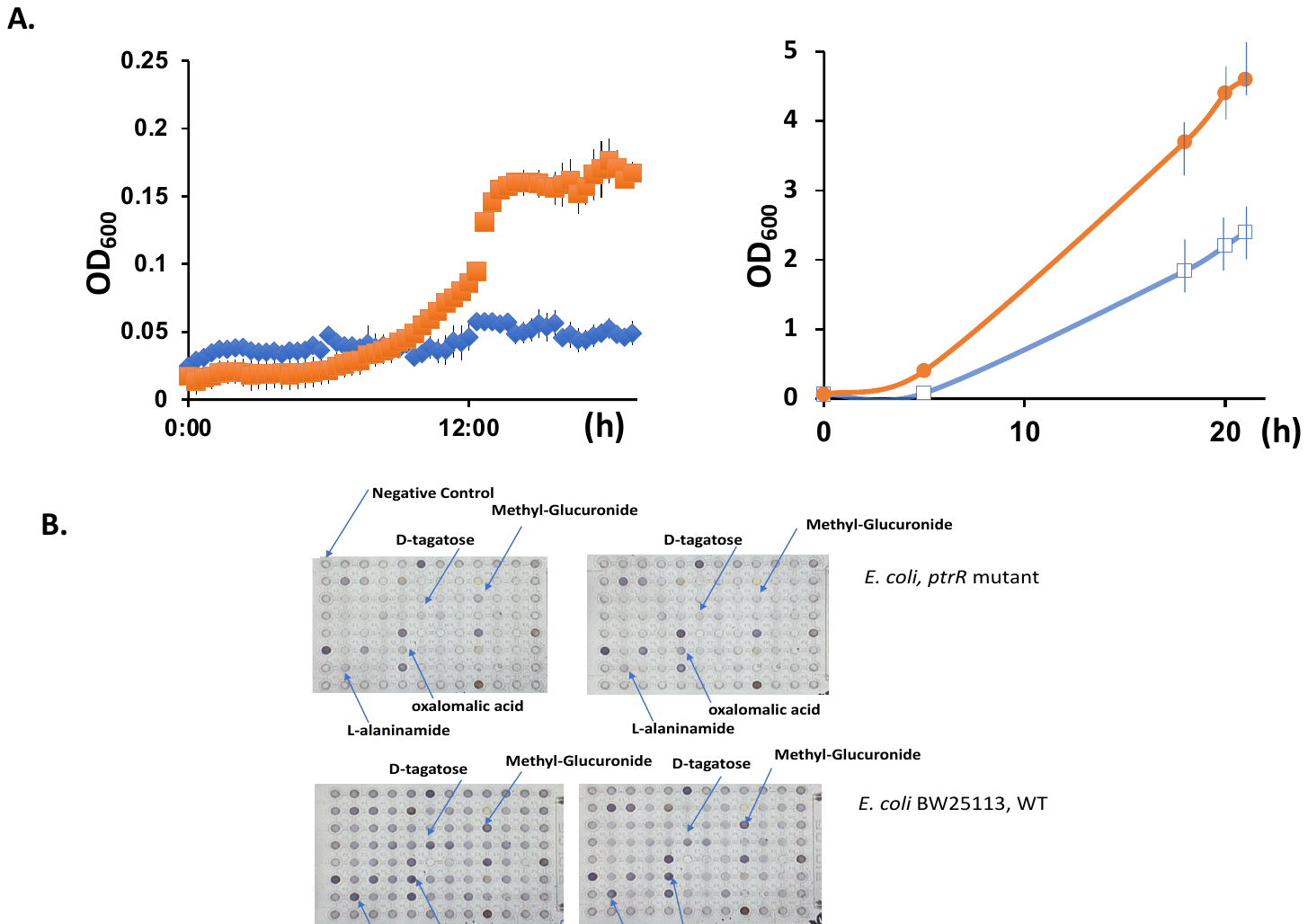

**Fig. S10** The antibiotic resistance phenotype for the *ptrR* (*yneJ*) deletion mutant and *E. coli* BW25113 wild type (blue line) for the plate PM11C, Biolog for two different concentrations of tetracycline is **A.** 1X and **B.** 8X.
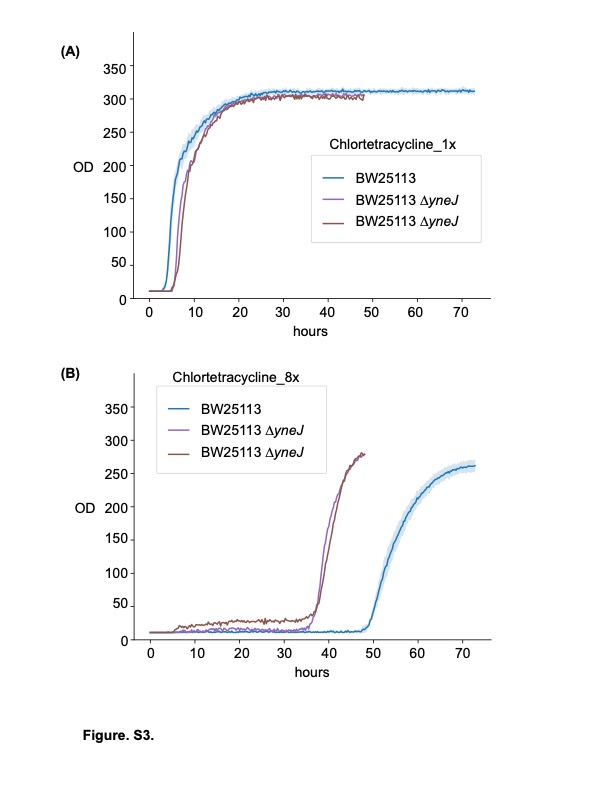

**TABLES**

**Table S1**

| b-number | *gene* | | Sequence | Position | Score | Log^2^ fold change |
| --- | --- | --- | --- | --- | --- | --- |
| b2724 | *hycB* | | GccGGACAAAGAACTC | -447 | 5.65 | -9.0 |
| b0903 | *pflB* | | GATGGACAAAGCGTTC | -356 | 7.26 | -5.39 |
| b1200 | *dhaK* | | GATGCACAAtGGATTC | -92 | 6.22 | -1.39 |
| b1241 | *adhB* | | GATGCtgAAAGGTGTC | -274 | 5.16 | -5.70 |
| b2503 | *pdeF* | GActCACAAAaGGTTC | | -365 | 5.16 | - |
| b4076 | *nrfA* | GATGaAtAAAGGGCTt | | -390 | 5.37 | - |
| b4020 | *yjbB* | GATtCACAAAtCTGTC | | -230 | 5.36 | - |
| b4028 | *yjbG* | GcTGGACgAAGAAGTC | | -153 | 4.80 | - |
| b1468 | *narZ* | GATGtACAAccGTGTC | | -75 | 5.87 | -2.28 |

**Table S2**
